# Cell Geometry and Junction Arrangement Define the Mechanical Robustness of Plant Tissues

**DOI:** 10.64898/2026.09.07.749888

**Authors:** Euan T. Smithers, Mahwish Ejaz, Martin Lenz, Leo Serra, Elise Laruelle, Sarah Robinson

## Abstract

Plant tissues are composed of immobile, pressurized cells that must maintain structural integrity under diverse environmental loads. Unlike animal tissues, which adapt through cellular rearrangement, plants must achieve mechanical robustness through the geometric configuration of their cellular networks. In this study, we establish a mechanistic link between cell geometry and mechanical resilience by integrating multilayer 3D mechanical simulations with cellular-resolution image analysis across a phylogenetically diverse panel of plant species, including *Zea mays*, *Tradescantia zebrina*, and *Arabidopsis thaliana*. We identify three-way junctions as fundamental mechanical elements that function as flexible hinges enabling mechanical strain accommodation in plant tissues. In contrast, four-way junctions are significantly stiffer and lack this strain-absorbing mechanism, providing a mechanical rationale for their biological avoidance. We also find that tissue material properties and strain response depend on edge lengths, cell layer, the degree of hexagonal shape, and turgor pressure response. These findings reveal the mechanism by which cell division patterns can actively tune tissue resilience to mechanical stress. This work provides new insights into the evolution of different cell shapes and offers clear principles for bio-inspired material science and tissue engineering.

**Summary for a general scientific audience:** Unlike animal cells, which can rearrange to accommodate mechanical stress, plant cells are encased in rigid walls and fixed within an immobile architecture. While much is known about the molecular composition of the cell wall, the mechanical consequences of the 3D cellular arrangement itself remain poorly understood. Here, we combine multiscale 3D simulations with high-resolution imaging to demonstrate how cell geometry and junction topology, specifically the prevalence of three-way versus four-way junctions, dictate tissue-level robustness. We show that specific packing arrangements minimize mechanical strain and explain why certain architectures are prevalent across diverse species.evolutionarily conserved. Our findings provide a quantitative framework for understanding how tissue topology, independent of wall biochemistry, serves as a primary determinant of plant structural integrity.

## 1 Introduction

Multicellular organisms face a fundamental mechanical challenge to build robust and functional tissues from individual cells that must withstand diverse and unpredictable environmental loads. In animal tissues, structural adaptation during development is facilitated through active cell migration and neighbour exchange [Walck-Shannon and Hardin, 2014, Alt et al., 2017]. Plant tissues contrast this dynamic behaviour as they are composed of immobile and pressurised cells encased in rigid interconnected walls that permanently fix tissue architecture [Cosgrove, 2005, Ali et al., 2023]. Plant tissue architecture must also be strong enough to endure continuous external mechanical stresses such as wind, gravity, insect landing, and soil resistance, stiff enough to avoid fracture, yet light and economical in their use of biomass [Moulia et al., 2021, Ashby and Medalist, 1983]. Plants must therefore achieve mechanical robustness during deformation while maintaining efficiency and developmental control within their fixed cellular networks.

Plant cells display a remarkable diversity of shapes ranging from complex lobed pavement cells and elaborate trichomes [Vőfély et al., 2019, Mathur, 2004] to the more uniform epidermal arrangements found in many species. Even within the epidermis the architecture varies significantly across plant lineages, including the hexagonal honeycomb structures in *Tradescantia zebrina* leaves and *Marchantia polymorpha* gemmae and the rectangular cell files in *Zea mays* leaves and *Arabidopsis thaliana* hypocotyls alongside the square arrangements in mosses such as *Atrichum crispum*(Fig. 1 and 6). This morphological variation prompts a fundamental inquiry regarding which mechanical drivers govern this diversity of shapes and how these distinct geometries impact the mechanical response of pressurised tissues.

**Figure 1:**
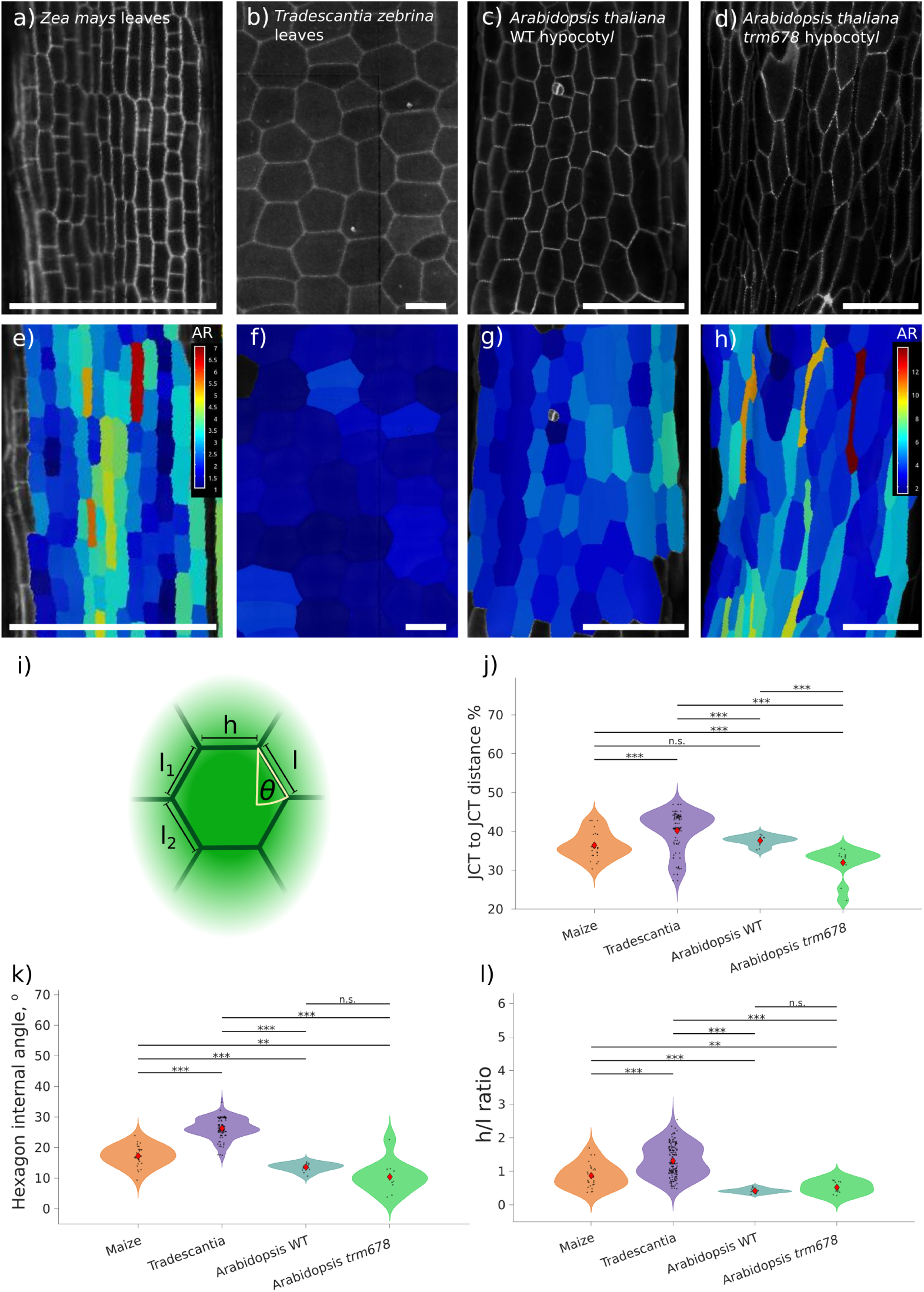
Quantification of cell shape and arrangement in different plant species. Confocal image of a) *Zea mays* leaf, b) *Tradescantia zebrina* leaf, c) Arabidopsis wild-type hypocotyl, and d) Arabidopsis *trm678* hypocotyl. e-h) The cell aspect ratios of the different species with e-g) sharing the same scale bar and h) on its own scale. e) *Zea mays*, f) *Tradescantia zebrina*, g) Arabidopsis wild type, h) Arabidopsis *trm678*. a-h) Scale bar is 100*µm*. i) Idealised hexagonal cells and the extracted metrics: the edge lengths *l*_1_,*l*_2_,*l*, *h* and the internal angle *θ*. j-l) Maize, *Tradescantia*, Arabidopsis wild type and Arabidopsis *trm678*. Each data point is the median value (from all the cells) from each independent sample (Maize *n* = 27, *Tradescantia n* = 100, Arabidopsis wild type *n* = 13 and Arabidopsis *trm678 n* = 12). By Welch’s ANOVA test, the means are significantly different (p*<* 0.001), and post hoc test by Games-Howell, = *p <* 0.05, = *p <* 0.01, = *p <* 0.001 and ns = not significant. j) The junction-to-junction distance as a percentage. k) Hexagon internal angle. l) Ratio of the transverse wall length (h) to longitudinal wall length.

While wall properties have long been known to influence tissue mechanics [Smithers et al., 2024, Cosgrove, 2005, Peaucelle et al., 2012, Hamant and Traas, 2010, Chen et al., 2025], we still lack a systematic, quantitative understanding of how cell geometry, arrangement, and junction topology integrate to define the mechanical response of turgid, multicellular plant tissues. Several studies have indicated that cell geometry and arrangement modulate tissue mechanical responses. For instance, cell file orientation can induce mechanical anisotropy [Majda et al., 2022, Malek and Gibson, 2017, Shafayet Zamil et al., 2017], while the effective mechanical stiffness of a tissue is often dependent on the internal angle of hexagonal cells [Majda et al., 2022, Malek and Gibson, 2017, Ashby and Medalist, 1983, Gibson, 2012, 2005]. Furthermore, plants manage stress through specific geometric adaptations: growth into elongated shapes can relieve local stresses [Sapala et al., 2018], and varying cell size in inner layers helps alleviate epidermal stresses [Silveira et al., 2025]. Additionally, junction topology is critical; four-way junctions, which are typically avoided [Sinnott and Bloch, 1941], can create weak spots for airspace formation [Sinnott and Bloch, 1941, Zhang et al., 2021]. Finally, investigations into cellular geometry have demonstrated the importance of cell density, with higher values increasing tissue stiffness and thereby regulating overall tissue properties [Malek and Gibson, 2017, 2015]. Despite this body of work, we still lack a systematic, quantitative comparison of how cell shape, arrangement, and junction arrangement together determine the mechanical behaviour of turgid, multicellular, multilayered plant tissues. Prior studies [Majda et al., 2022, Malek and Gibson, 2017, Ashby and Medalist, 1983, Gibson, 2012, 2005, Lee et al., 2025, Shafayet Zamil et al., 2017] have typically varied one geometric parameter at a time, often in idealised, single-layer models, or without accounting for turgor, and without a detailed cellular level comparison with experimental data, leaving us without a complete overview.

We address this knowledge gap here by integrating multilayer 3D mechanical simulations with quantitative cellular resolution experimental analysis. By systematically varying cell shapes and junction topologies we demonstrate that tissue deformability is tunable through cellular geometry alone. We identify geometries that minimise stress, explain why tissues with predominantly three-way junctions deform differently from those containing four-way junctions and finally provide a quantitative mechanistic framework linking tissue architecture to mechanical resilience.

## 2 Results

### 2.1 Quantification of cell geometry in different species

To study the effect of cell shape and arrangement on tissue mechanical properties, we selected a phylogenetically diverse panel of tissues exhibiting varied cell geometries. We identified *Zea mays* (Maize) undifferentiated leaf primordia with rectangular cells (Fig. 1a, e), *Tradescantia zebrina* (*Tradescantia*) leaf epidermis with cells that are near perfect hexagons (Fig. 1b, f), and *Arabidopsis thaliana* (Arabidopsis) hypocotyls to investigate cell shape in a 3D rectangular organ model (Fig. 1c, g). In order to quantify how the different species vary, we collected confocal images and quantified the geometric properties of the tissues using metrics from existing literature, namely the aspect ratio, junction-to-junction distance [Sinnott and Bloch, 1941], hexagon internal angle and edge length ratio [Gibson, 1989] (Fig. 1i). Briefly, cell junctions are identified as vertices where three or more walls intersect.

To quantify the relative junction position along the cell lengths, we identified the two longitudinal edge segments (*l*_1_ and *l*_2_) split by each junction along the cell file (Fig. 1i; Section 9.2.2) and calculated a percentage as 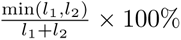 which we defined as the junction-to-junction distance. Maize leaves had a junction-to-junction distance of 36% ± 0.03 SD, Tradescantia leaves the largest, with an average junction distance around 40%±0.05 SD, and wild-type Arabidopsis hypocotyls with 38%±0.01 SD (Fig. 1j). In addition, we also measured the junction distance of an Arabidopsis cell division mutant impaired in preprophase band formation, but does not impact interphase microtubules, TON1 Recruiting Motif *trm678* [Schaefer et al., 2017]. The *trm678* hypocotyls displayed defects in division plane orientation (Fig. 1d, h), resulting in a reduced junction-to-junction distance of 32% ± 0.04 SD, showing how cell division defects can alter the cellular arrangement and noise.

Hexagon internal angle is defined as the angle the longitudinal walls are oriented away from the perpendicular direction relative to the transverse wall they are connected to [Gibson, 1989] and was calculated in all species (Fig. 1k) (see section 9). For Tradescantia, internal angles clustered around a mean value of 26.3*^◦^* 3.2*^◦^* SD confirming symmetric well ordered hexagonal cell packing (a regular hexagon has an angle of *θ* = 30*^◦^*). In contrast, Maize cells had an internal angle of 17.3*^◦^* 3.4*^◦^* SD, as longitudinal walls align more orthogonally. The internal hexagonal angle was further significantly lower in Arabidopsis wild-type hypocotyls (13.6*^◦^* 1.3*^◦^* SD). Finally, the smallest internal angle was in the hypocotyls of *trm678* mutants, with an internal angle of 10.4*^◦^* 5.1*^◦^* SD.

To further quantify the variation in geometry, we calculated a ratio of the transverse wall length (h) to longitudinal wall length (l) [Gibson, 1989, Boudaoud et al., 2023] (h/l) for individual cells of each species. The h/l ratio can measure the cell aspect ratio, providing a quantitative metric of cell geometry along their principal axis. Tradescantia leaf epidermal cells show an average h/l ratio of 1.3 ± 0.5 SD, representing more isotropic cell geometry. In contrast, Maize cells showed an h/l ratio of 0.85 ± 0.4 SD, wild-type Arabidopsis with a ratio of 0.4 ± 0.07 SD and *trm678* with 0.5 ± 0.2 SD (Fig. 1l).

We have identified three key geometric metrics that have quantifiable differences between species for further theoretical investigation

### 2.2 Cell geometry alters stress distribution

Understanding the impact of cell shape is vital for determining how tissues manage turgor-induced stresses. This factor is particularly pertinent for epidermal cells, which face elevated stress due to the absence of adjacent cells on their outer surfaces. Because larger cell dimensions experience higher cell stress [Sapala et al., 2018], identifying a cell geometry that efficiently tessellates an area with minimal cell stress while utilising the same amount of resources is of great interest.

To evaluate this, we established a fixed area to be tiled and a fixed available perimeter, representing the resource constraints across various shapes, and subsequently calculated the number of cells that could accommodate this space. Using this cell count, we identified the largest empty circle capable of fitting within the cell boundaries (see section 9.1.1 for details). This radius serves as an approximation for cell stress, given the positive correlation between the two parameters [Sapala et al., 2018].

Given identical resource limits, we found that a greater number of hexagonal cells can be generated compared to squares and rectangular ones (Fig. 2a), consistent with earlier research showing hexagons have an exceptionally efficient configuration for spatial packing [Hales, 2001]. Nevertheless, an evaluation of the maximum internal radius revealed that hexagons and squares align along an identical curve, whereas rectangles display a smaller radius (Fig. 2b). We verified that this radius effectively approximates stress by simulating the inflation of these geometries through finite element analysis (Fig. 7a). This confirmed that smaller cellular volumes experience reduced stress levels, and that the maximum internal radius reliably correlates with stress across diverse cell geometries, corroborating foundational observations.

**Figure 2:**
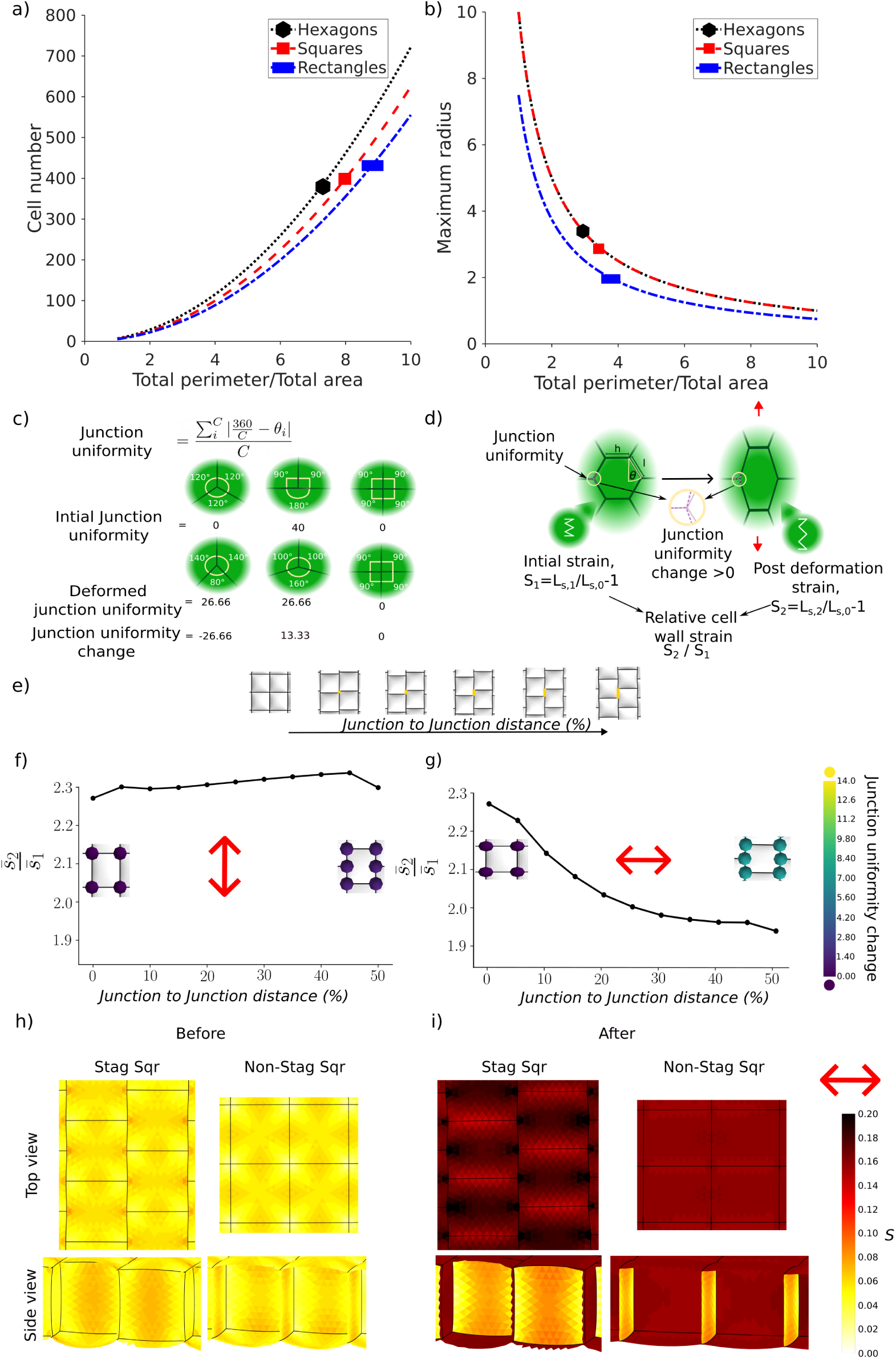
The impact of cell shape and arrangement on wall strain. a-b) Comparison for different amounts of resources (the total allowed perimeter) of squares (red dashed line), rectangular (blue dashed dotted line), and hexagonal cells (dotted line) of a) the number that fill an area with a set amount of resources and b) the radius of the largest circle we can fit into the shapes. c) The junction uniformity definition. Defined as the average degree to which an angle differs from the uniform angle at a junction. d) Diagram depicting how three-way junctions can change their junction uniformity to allow shape change during deformation (represented as the red arrows) and the relative wall strain measure. Also shown are the labels *h* and *l* for the different side lengths, the internal angle *θ*, and the principles behind the strain ratio. e) Square cell outlines as the junction-to-junction distance is increased from four-way junctions to perfectly staggered three-way junctions (yellow). f-g) The average strain ratio (the weighted average of each triangle’s principal strain in the mesh after deformation divided by the same measure before) when deformed by 20% in the f) cell file direction and g) perpendicular to the cell file direction while varying the junction-to-junction distance along the cell edge length, with the distance as a percentage along the cell’s side length. The cell outlines post-deformation are also overlaid on the figure, with the coloured dots indicating the average junction uniformity change in the tissue. h-i) The principal strain shown in the staggered and non-staggered square setups from the top and side view of a cut mesh before and after deformation perpendicular to the cell file.

These results build upon those of Sapala et al. [2018], showing that rectangles are not only advantageous in a single-cell context but also in a tissue context.

### 2.3 Three-way junctions function as hinge mechanisms to accommodate mechanical strain

To investigate the consequences of different cellular arrangements and geometry on tissue properties, we created meshes of different idealised shapes. We varied junction-to-junction distance, the hexagon internal angle, and wall length ratios to capture the variation observed in the biological species (Fig. 1). The idealised meshes were inflated to mimic the actions of turgor pressure, then subjected to a deformation up to a maximum of 20% in the longitudinal direction with the cell file and the transverse directions against the cell file.

To analyse the simulations, we introduced a metric called junction angle uniformity, which evaluates shape changes by measuring how evenly the walls or wall angles are distributed around a junction, 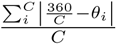 (Fig. 2c). For example, if the walls are equally spaced around the junction, they all have the same angle and the junction angle uniformity is zero. We then examined the junction uniformity change by calculating the difference in junction angle uniformity before and after deformation (similar to [Wang et al., 2024b]). We also compared the strain on the cells, which we calculated as the cell wall’s average strain (across all walls) after deformation divided by the average strain of the inflated shape before deformation (Fig. 2d). This normalises for the different initial strains between the meshes with different shapes.

#### 2.3.1 Junction-to-junction distance determines junction deformability

In plants new cell walls tend to be positioned more than 10% of the total wall length away to avoid creating four-way junctions [Sinnott and Bloch, 1941]. Four-way junctions seem to be actively avoided [Martinez et al., 2018, Wang et al., 2024b] in a mechanism that may be guided by stress minima in the cell wall at existing three-way junctions [Gascon et al., 2025]. Four-way junctions do occur in some plant species (*Marchantia*, [Bonfanti et al., 2023] and a range of mosses, 6), potentially associated with thicker cell walls [Sinnott and Bloch, 1941], wound-induced division [Lloyd, 1991] or air space formation [Sinnott and Bloch, 1941, Zhang et al., 2021]. The consequences of avoiding four-way junctions in plants are not known.

To investigate the impact of having four-way junctions, we created ideal meshes with different distances between the junctions ranging from 0% for four-way junctions to 50% for perfectly spaced three-way junctions (Fig. 2e). When we examined the strain ratio for the different setups, at the extremes, four-way junctions and staggered three-way junctions (0% and 50% junction-to-junction distance, respectively) displayed little difference in the strain ratio when deformed in the cell file direction (Fig. 2f). When deformed orthogonally to the cell file (Fig. 2g), the arrangements with the larger junction distance (staggered) exhibited a smaller increase in strain compared to the four-way junction versions (non-staggered) (matching the lower transverse stiffness previously observed [Majda et al., 2022, Malek and Gibson, 2017, Shafayet Zamil et al., 2017]). Setups with a junction-to-junction distance within 10% of their side length experienced a strain increase on par with four-way junctions, perhaps explaining the observed absence of walls within 10% of existing junctions rather than only four-way junctions being avoided.

To understand the differences in the strain ratios, we analysed the junction uniformity changes. When deformed against the cell file direction, the staggered setup’s junction uniformity changed (the coloured nodes in Fig. 2g). There is, therefore, a relationship between three-way junction setups allowing a junction uniformity change and a reduction in the strain upon deformation, which is absent when deformed in the cell file direction (Fig, 2f-g). The three-way junction structure thus act as hinges, a property important for energy absorption [Ashby and Medalist, 1983], and thus allows a shape change, meaning the tissue is deformed without the edges being extended (Fig. 2d).

In animal tissues, three-way junctions are recognised as mechanically sensitive locations with tight regulation [Higashi and Miller, 2017, Bosveld et al., 2016, Yu and Zallen, 2020], four-way junctions are predicted to open more easily for epithelial integration [Ventura et al., 2022]. To understand the impact of the junction arrangement on the distribution of strain in our tissues, we examined the strain across the full 3D shape (Fig. 2h, i). We observed that the three-way junction hinge reduced strain in the anticlinal walls along the stretch directions. A strain hotspot is obvious at the junction in the three-way junction setups (Fig. 2h, i); this is true whether the initial junction had a maximum angle of 120*^◦^* or 180*^◦^* . To examine this hotspot further, we examined the maximum strain ratio (as the average will not capture the strain hotspot) on the top surface before and after deformation (Fig. 7d-e). The three-way junction setups exhibited a reduced increase in the maximum strain ratio, perhaps making them less vulnerable to perturbations if they have already been reinforced.

Four-way and close three-way junctions (*<* 10%) are thus more vulnerable to greater strain increases during deformation due to their lack of flexibility. This is consistent with the observations from the animal literature.

#### 2.3.2 Tissue experiences greater transverse or longitudinal strain dependant on the junction angle

The internal angle of the cells varied between the species we investigated to give rectangular or hexagonal cells. To identify the impact on tissue mechanical properties we generated idealised meshes with altered internal angles while controlling for the side length to reflect the differences we saw in our chosen study species (Fig. 3a, and Fig. 1k). Upon increasing the angle, cells had lower strain in the cell-file direction (Fig. 3b). This was reversed when the strain was applied transversely (Fig. 3c). This result is in agreement with previous analytical work [Gibson, 2005, Malek and Gibson, 2015, Gibson, 1989] and [Shafayet Zamil et al., 2017], which examined the hexagon aspect ratio. These previous works were not performed under turgid conditions, meaning we have now confirmed the same angle-dependent relationships under turgid conditions. The perfect regular hexagonal shape (30*^◦^* internal angle) has a strain ratio value that lies between the maximum and minimim values in both deformation directions, demonstrating its versatility. The degree of junction pinch in from a rectangle to a compacted hexagon thus affects the tissue’s wall strain.

**Figure 3:**
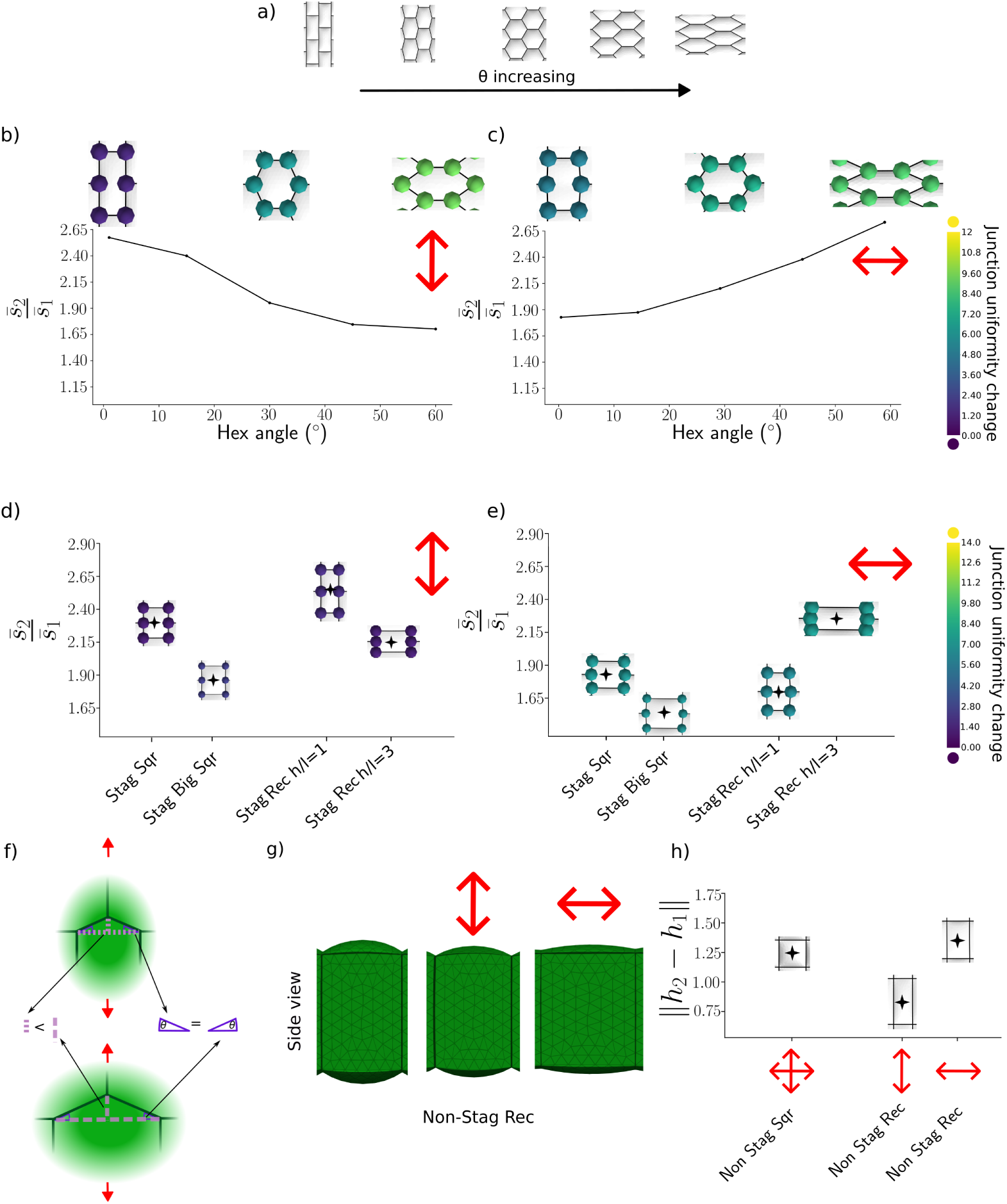
Junction angles and edge length also affect how cells respond to deformation. a) The different hexagonal cell outlines with increasing internal angle, all having the same top surface area and equal sides. b-c) The average strain ratio as the hexagonal internal angle changes when deforming by 20% in the e) cell file direction and f) perpendicular to the cell file, overlaid with the outlines of the hexagonal setup post-deformation, with the coloured dots showing the average junction uniformity change in the tissue for *θ* = 0, 30, 60. d-e) The average strain ratio comparing staggered setups, specifically staggered squares with side length 10*µm* and their larger counterpart with side length 15*µm* and staggered rectangles with *h/l* = 1 and staggered rectangles with *h/l* = 3, when stretched by 20% in the d) cell file direction and e) perpendicular to the cell file direction. The star indicates the value on the strain-ratio scale, and the outlines of the different post-deformation setups are overlaid, with the coloured dots showing the average junction uniformity change. f) Diagram showing the effect of a longer edge (on the bottom) upon deformation (red arrows). For the same angle *θ* in both setups, the shape change (the dashed line’s length) is greater in the bigger square. g) Side view of the cut meshes of the non-staggered rectangles before and after deformation in the two different directions, showing different epidermal wall bulging heights. h) The absolute height difference between before and after deformation between the non-staggered squares and non-staggered rectangles meshes deformed in both directions, with the star showing the value of the height difference.

#### 2.3.3 Wall length impacts tissue properties

Our selected study species (Fig. 1l) also varied in wall length (h/l ratio). We investigated the consequences of varying cell wall length by comparing squares of different sizes and rectangles of different aspect ratios. We found that smaller squares had a higher strain increase when deformed in either direction (Fig. 3d, e). For the rectangles, the strain was highest when they were deformed along their longest axis (Fig. 3d, e). Deforming transversely caused a similar junction uniformity change for the different shapes, but the strain ratio increase was lower in the shapes with a longer wall perpendicular to the deformation direction (Fig. 3e). For a similar junction uniformity change, a longer edge allowed more shape change (Fig. 3f), resulting in a lower strain ratio.

Another factor is how much the periclinal wall bulging is allowed to change (Fig. 3g, h). This effect can be observed when comparing the staggered rectangles of different aspect ratios, as well as staggered squares and rectangles (and non-staggered Fig. 7h). The periclinal wall of the non-staggered rectangles bulged out by different amounts after being pulled in different directions, with the height changing more when pulled against the cell file (Fig. 3g, h). The height change difference between squares and rectangles also explains why the square strain ratio increases less than the rectangles when deformed in the cell file direction (Fig. 3d). How restricted the height changes are, affects the structure’s flexibility and may help reduce strain on the periclinal wall upon deformation. Thus, both the cell edge length and the extent of periclinal wall bulging affect the deformation response (explaining similar observations made by [Majda et al., 2022]). Note that possible differential periclinal wall bulging is not the reason for the differences in three-way and four-way junction deformation response. If this were true, we would see more differences when the different junction-to-junction distance square arrays are deformed in the cell file direction (Fig. 2f). Thus, both junction uniformity change and other geometric factors affect cellular response to deformation. To confirm that our results are not an artefact of using the strain ratio as a metric, we verified that the result is unchanged when the strain difference is considered (Fig. 8a-b). Furthermore, this is also not dependent on the turgor pressure implementation, where with non-constant turgor (Fig. 7b-c), we obtained the same results.

Many plant cells have anisotropic cell walls [Baskin, 2005] due to the presence of aligned cellulose fibres, meaning their stiffness varies with the direction of deformation. Simulations were therefore repeated with anisotropic periclinal walls, with the anisotropy and deformation direction specified perpendicular to the cell file. Similarly to the above, three-way junctions showed a reduced strain ratio (Fig. 7f), albeit with a lower junction uniformity change than in the non-anisotropic case. Thus, anisotropic cell walls do not change our conclusions.

In a monolayer, we have thus shown that plants can modulate the degree of strain a tissue experiences upon deformation by altering junction-to-junction distance, internal angle, and edge length.

### 2.4 Geometric Determinants of Strain Persist in Multilayered Tissues

To further investigate the consequences of cell size and wall length on 3D tissue-scale properties, we developed multilayer simulations. Idealised meshes were generated with an inner cortex layer and an outer epidermal layer on the top and bottom (Fig. 4a). These simulations enabled us to investigate our predictions in a more realistic context, as plants are multilayered structures with internal layers influencing tissue mechanics [Silveira et al., 2025].

**Figure 4:**
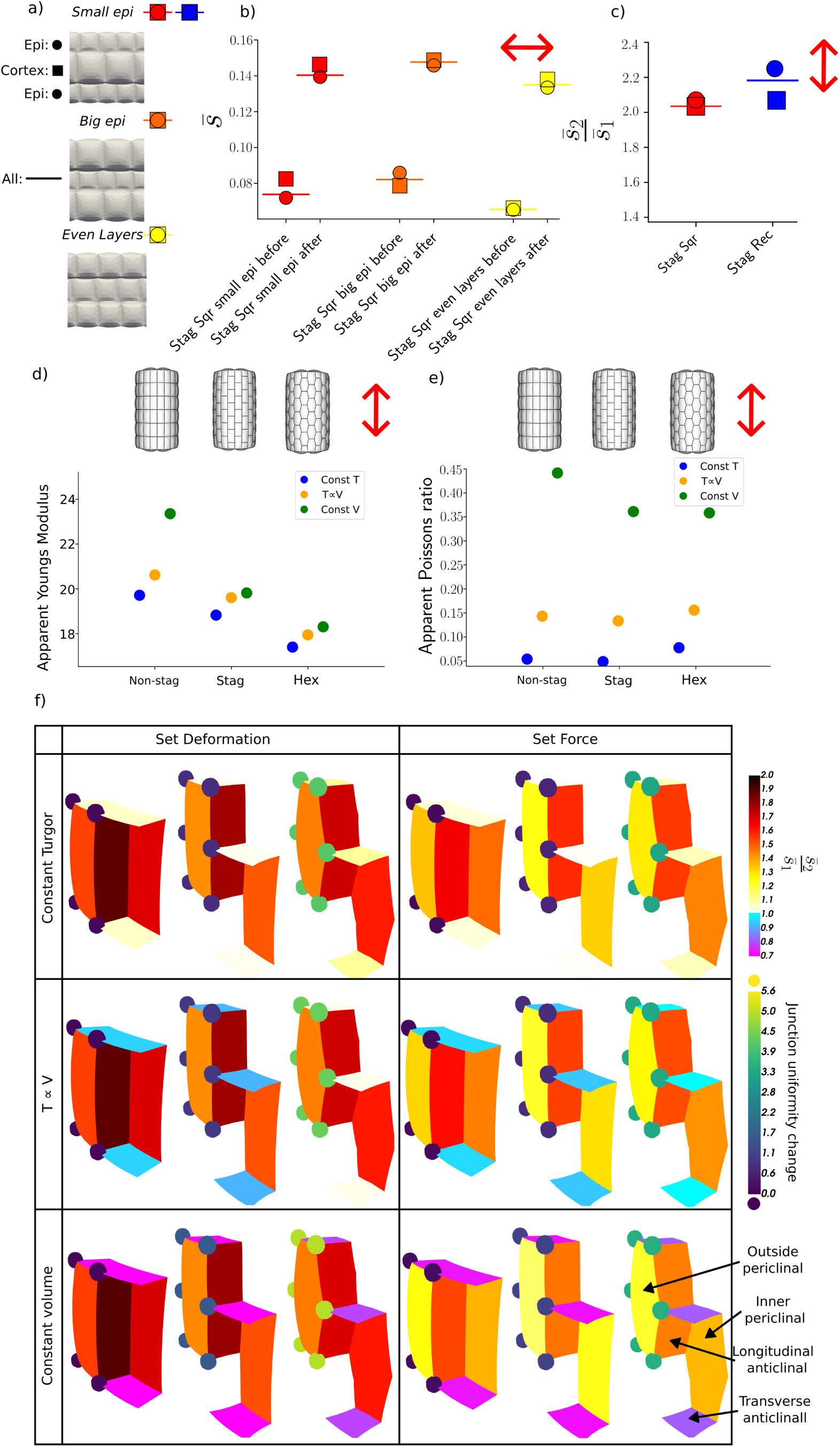
Multilayer simulations of deforming tissues. a) Side view of the three multilayered variants, one with a small epidermis (staggered squares, red and rectangles, blue) where each cell on the epidermis has total top surface of 100*µm*^2^ and with the cortex’s having a value of 225*µm*^2^), another with a big epidermis (staggered squares, orange with the size in the layers reversed) and even layer variant (staggered squares, yellow, all of the smaller size). The average strain of all layers is depicted as a line, the epidermal values as a circle and the cortex as a square. b) The average strain of staggered square with the different size variants, small epidermis (red), big epidermis (orange), and even layers (yellow), before and after deformation when deformed by 20% perpendicular to the cell file direction c) The average strain ratio comparing staggered squares (red) and rectangles (blue) when deformed by 20% in the cell file direction. d-e) Material properties of cylindrical multilayered non-staggered, staggered and hexagonal setups calculated by applying a set force axially on the top and bottom in the different turgor regimes: constant turgor (blue), turgor is linearly proportional to the volume (yellow) and constant volume (green). d) The apparent Young’s modulus. e) The apparent Poisson ratio. f) The average strain ratio in the non-staggered, staggered and hexagonal setups upon deformation on the periclinal outside and inside wall, and the anticlinal axial and transverse walls (labelled with the arrows) for the different turgor regimes. The dots show the average junction uniformity change in the tissue.

Typically, the inner layers of plant tissues have larger cells than in the outer epidermal layers. To determine the extent to which cell geometry versus layer position affects the strain response, we generated idealised meshes with small epidermal cells, larger epidermal cells or in which all layers were of equal size (Fig. 4a). The simulation where all cell layers were the same size revealed that the inner cortex strain increased more when deformed transversely (Fig. 4b) (not observed under longitudinal stretching, Fig. 7i), a factor that exists in the other multilayer geometries. This is likely because the cortex is sandwiched between the other layers and consequently exposed to more junctions: its own layers and those of the two epidermal layers, which are strain hotspot locations. Strain increase is higher for smaller cells irrespective of their position in the tissue, confirming the relationship with edge length (Fig. 4b and Fig. 7i, j). The cellular bulging effect can be observed in comparing the rectangular and square setups deformed in the cell file direction, where the strain ratio in the epidermis differs, but the cortex cells do not (the cortex cells do not bulge out) (Fig. 4c). When deforming against the cell file, the cortex in three-way junction setups still showed a reduced strain increase compared to four-way junctions (Fig. 7k), indicating that junction uniformity changes are a dominant factor in deformation response. These multilayer simulations confirm that the cellular geometric control over tissue mechanics is not limited to simple layers, but functions as a robust regulatory mechanism even within the heterogeneous cellular environment of internal tissue layers. The simulations highlight the vulnerability of internal layers to strain hotspots, and the critical role they play in mediating strain distribution across the tissue.

### 2.5 Cell geometry has a stronger influence in cylindrical structures

To test our findings in other tissue geometries, we examine cylindrical structures more like those seen in hypocotyls. This also enables us to investigate the impact of the additional tissue-scale hoop stress typically seen in cylindrical shapes [McKeen, 2016]. Because these different arrangements have the same top surface areas, we can determine their tissue-scale material properties in a controlled manner. The four-way junctions are the stiffest, followed by the staggered setups and then the hexagonal setups (Fig. 4d) for all turgor regimes. These results differ from the monolayer simulations (Fig. 7g and previous monolayer simulations [Malek and Gibson, 2017, Majda et al., 2022]) where, in the cell-file direction, the stiffnesses of the staggered and non-staggered arrangements were less pronounced. This difference from the monolayer simulations arises from the geometry’s cylinder-like structure, demonstrating the importance of replicating actual geometries when simulating deformed tissues. However, these results agree with the observation that, for the same cell number, hexagonal structures are softer [Malek and Gibson, 2017]. We wanted to further understand the role of turgor in the model and the potential effect on deformation. Specifically, how the method of implementing turgor impacts the outcome. We therefore repeated simulations using either constant volume [Diels et al., 2019, Weber et al., 2015] where no water leaks during deformation; constant pressure [Weber et al., 2015], whereby water is taken up at a steady rate to maintain constant turgor pressure; or pressure changes proportional to volume changes (*T ∝ V*) [Vesenjak et al., 2007] such that water behaves like a gas.

We see that the Young’s modulus, which is a measure of the elastic properties of the tissue, is highest (stiffer) for constant volume (Fig. 4d), constant turgor sits at the other extreme, with the linearly proportional turgor model providing in-between results. The difference in Poisson ratio, which is a measure of the deformation of the tissue perpendicular to the direction of the applied load, between the different geometries is also affected by turgor implementation (Fig. 4e). In the case of constant turgor and *T ∝ V*, it slightly increases in the hexagonal setup, and in the case of constant volume, it increases for non-staggered, being close to 0.5, which is expected due to water incompressibility. We have therefore shown that overall tissue properties are severely affected by what is happening to the water during deformation. However, the impact of shape qualitatively remains the same (Fig. 4f).

In the hypocotyl model, we can see in detail the strain ratio on the different walls (Fig. 4f). The longitudinal anticlinal walls have the largest difference in strain ratio in the non-staggered, staggered, and hexagonal setups, with the outside periclinal wall exhibiting a smaller difference. We also see that the tangential anticlinal walls are relatively compressed, particularly in the constant-volume turgor case. The hexagonal structures exhibit less relative compression, demonstrating that the flexible structure helps relieve both tension in the longitudinal walls and compression in the transverse walls. Additionally, the inside periclinal wall has a higher strain ratio in all cases, possibly due to exposure to more junctions, which have strain hotspots.

In all these cases, again the three-way junction setup allowed a greater junction uniformity change and a reduced strain ratio increase upon deformation compared to the four-way junction counterparts. This difference increases further for the hexagonal shapes in the hypocotyl-like structure.

In conclusion, three-way junctions help a tissue reduce its strain upon deformation, with hexagonal shapes permitting this reduction from both directions. However, this creates softer tissues, and the three-way junction cannot be close to another junction. Other geometric factors influence this relationship, including the 3D tissue geometry, edge length, and epidermal bulging with turgor pressure also having an impact.

### 2.6 Junction change reduces strain in cell walls of living tissues when deformed

In live specimens we tested the theoretical predictions on the impact of junction distance, junction angle and cell length on the overall tissue properties and the strain distribution in the cells. Previously, differences have been observed in the stress-strain curves of cell arrays deformed with and against the cell file previously categorised [Majda et al., 2022, Shafayet Zamil et al., 2017], here we focus on how the geometries change upon deformation on a cellular level. An updated version of an automated confocal micro-extensometer was used to apply axial tension to species with different cellular geometries (Fig. 8f) [Robinson et al., 2017]. The samples were deformed by 5 − 20% and confocal images were obtained. All cells were segmented in their deformed and undeformed states (see section 9). The junction-to-junction edge lengths and junction changes were calculated before and after deformation. These before-and-after measurements were then mapped onto one another, allowing comparison of their changes to calculate the change in length 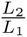 and the junction uniformity change (Fig. 8f).

To examine the edge strain and enable comparison with simulations, we characterised each edge into four categories based on the direction in which the tissue was stretched and whether the wall was aligned or orthogonal to this direction (Fig. 5a, section 9).

**Figure 5:**
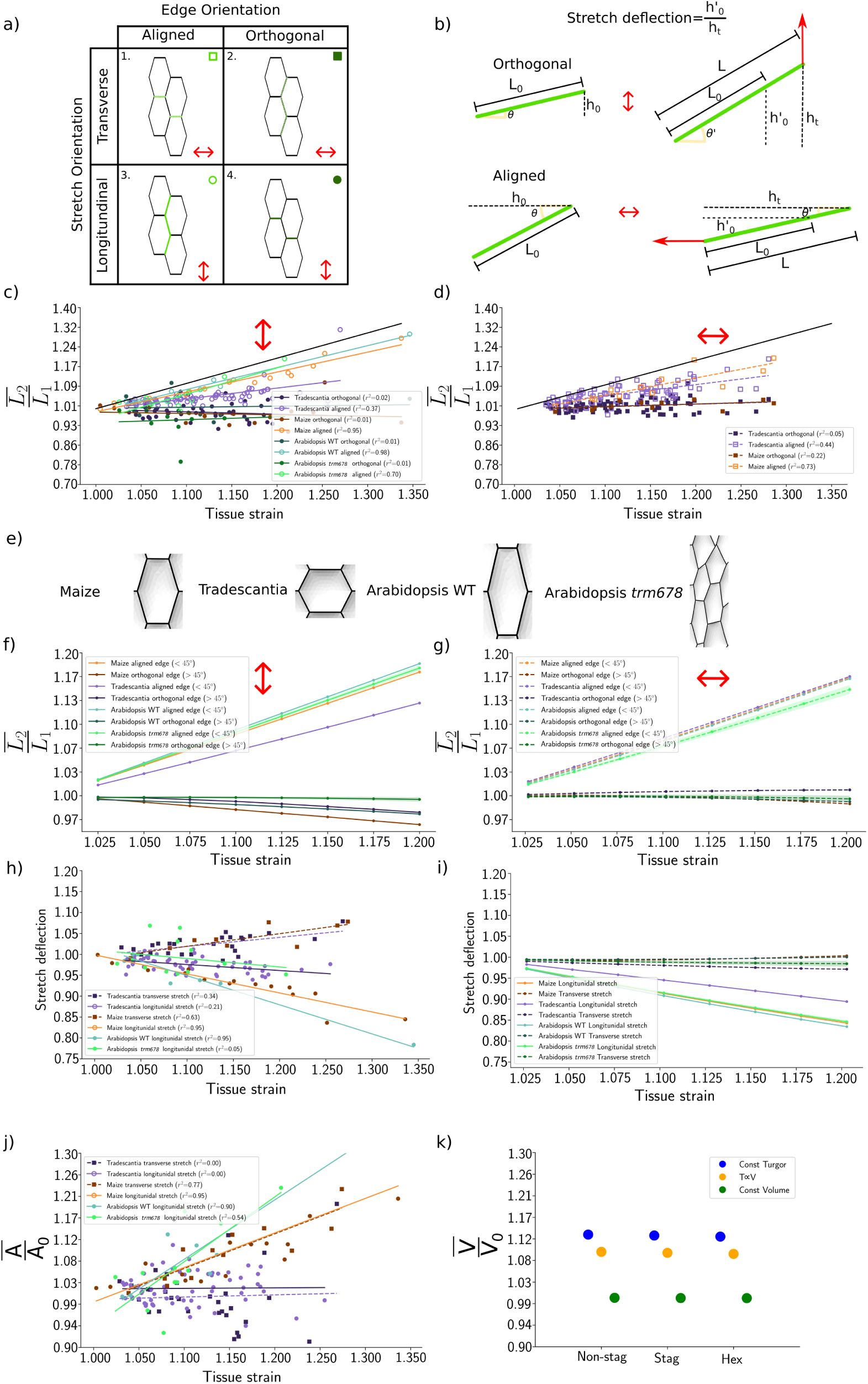
The deformation of live specimens and comparison with simulations. a) Edge-type classification, shown in green (solid for longitudinal, dashed for transverse) based on the direction the tissue was stretched in and if the wall aligned or was orthogonal to this direction, with a unique (green) dot style used in figures (square or circle, filled or unfilled, and light or dark) for each style. b) Diagram of stretch deflection for orthogonal and aligned edges. c-d) The edge length ratio of aligned and non-aligned edges as the tissue strain increases for *Tradescantia* (purple), Maize (orange), Arabidopsis wild type (blue) and Arabidopsis *trm678* (green) when deformed in the d) longitudinal and e) transverse direction. Data points represent sample medians (Maize *n* = 27, *Tradescantia n* = 100, WT *n* = 8, *trm678 n* = 12); lines show linear fits (*r*^2^ indicated); black line shows *y* = *x*. f) Simulation outlines per species (*trm678* shows 1 of 5). g-h) The edge length ratio of aligned and non-aligned edges as the tissue strain increases for the different species simulations when deformed in the g) longitudinal and h) transverse direction (green line/shading: *trm678* mean ± SD). i-j) Stretch deflection vs. tissue strain for all edges when deformed transversely or longitudinally (aligned and orthogonal edges grouped) in the i) experimental data and j) the simulated cells. k) The cell area changes during deformation in the longitudinal and transverse direction. l) Simulated cell volume change in cylindrical models across cellular arrangements and turgor regimes.

As tissue strain increased, the average edge length change per sample increased (*p <* 0.001 see section 9) for the aligned edges for all Maize, *Tradescantia*, Arabidopsis WT and Arabidopsis *trm678* (Fig. 5c-d). Additionally, edge Direction (*p <* 0.001), species (*p <* 0.001) and stretch Direction (*p <* 0.001) significantly determined the variance in edge strain response. Examining the different species’ response to longitudinal direction deformation (Fig. 5c), and comparing them against the *y* = *x* line, (the relationship of the edges exactly deforming with the tissue), showed the aligned edges of Arabidopsis WT to follow it more closely, followed by Maize and then *Tradescantia*. This acts in accordance with the behaviour expected with increasing internal angle of hexagonal structures (Fig. 3a-b). Note that, for example, if Maize were stiffer, this would scale all the data points down and thus would not change the relationship to the *y* = *x* line. On the other hand, the orthogonal edges show evidence of increasingly relative compression in all species as the tissue strain increases.

When comparing the tissues in the orthogonal direction (Fig. 5d) we see that Maize and *Tradescantia* show similar wall length changes (it was not possible to deform Arabidopsis hypocotyls transversely). Additionally, we see that the edge extension of Maize is less, and the *Tradescantia* is greater when compared to their longitudinal deformation, as predicted by our simulations (Fig. 3b-c) and previous results [Majda et al., 2022, Shafayet Zamil et al., 2017].

To further validate, for each species type we used the average *h/l* ratio and internal angle to create their average hexagonal shape (Fig. 5e). We then repeated our deformation simulations and now instead examined the change in length 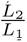 (Fig. 5f-g). These simulations agree qualitatively with the response observed in the data. In the longitudinal direction (Fig. 5f), Arabidopsis’s aligned edges have the highest edge extension, closely followed by Maize, and then *Tradescantia* with a gap in between. The orthogonal edges also all show relative compression, with the ordering between the species not as precisely following the data. In the transverse direction (Fig. 5g), we once again see agreement with Maize and *Tradescantia* being more similar in their response. We can also see the response of Arabidopsis if we were able to deform transversely, where it has a reduced edge extension. The orthogonal edges in this deformation all show less relative compression compared to longitudinal deformation, as also seen in the data.

To determine the extent to which junction angle change can explain the differences observed between species and stretch directions, we quantified the edge deflection (Fig. 5b). This represents the degree of displacement in the direction of deformation, driven by edge reorientation following junctional uniformity change. The edge deflection changed upon tissue deformation (*p <* 0.001 see section 9), species (*p <* 0.001) and stretch Direction (*p <* 0.001). When deformed in the longitudinal direction, this change was greatest in *Tradescantia* (Fig. 5h) (can also be seen in the junction uniformity change in Fig. 8d). The edge deflection decreased in the longitudinal direction as the junction changed as much as it could until the edge then had to take strain. In the transverse direction, we saw *Tradescantia* and, importantly, Maize had higher edge deflection, explaining why they exhibited less edge strain in this direction (Fig. 5h). Examining the edge deflection in our simulations, we saw the exact same pattern (Fig. 5i), further validating the predictive power of our model. We also observed continuous junction change in *Tradescantia*, whereas past work suggested that junctions changed before edge deformation, while assuming rigid edges [Masters and Evans, 1996, Gibson, 2005]. Thus, hexagonal cellular setups seen in *Tradescantia* exhibit a greater change in their junctions, as predicted by the modelling. There is, therefore, a relationship between the *Tradescantia* junctions changing more, due to their flexible hexagonal shapes and having a reduced strain upon deformation, as predicted by the simulations and past results.

Using the cell shape parameters extracted for all 6-sided cells (for Maize and *Tradescantia*, Fig. 8c; see section 9), Spearman correlation coefficients were calculated compared against the cell average edge deformations (Fig. 8c). Because tissue strain was the sole external driver, these moderate values indicate that other geometric factors also dictate edge-specific deformation. Specifically, *θ* in Maize and *Tradescantia* correlated negatively with edge strain along the cell file and positively across the cell, aligning with our simulations and past results [Gibson, 2005].

These moderate correlations stem from additional geometric noise. Hexagonal side length influences stiffness [Gibson, 2005], meaning concurrent changes in angle and edge length can offset each other—while array irregularities cause neighbouring cells with different internal angles to deform differently than expected. Together, these results demonstrate how local cellular geometry modulates individual cell deformation under tissue-level strain.

The analysis of experimental data demonstrates that our theoretical predictions hold true in live specimens. Therefore, geometric factors such as internal angles and three-way junctions influence tissue responses to deformation.

### 2.7 Axial loading causes an increase in cell area consistent with model of constant turgor pressure

We next explored the impact of the different simulated turgor regimes on cell area and the Poisson ratio to gain insight into the live-specimen properties. Walls which did not align with the direction of applied deformation showed a slight relative compression in both the experimental data (Fig. 5c-d) and in the model (Fig. 4f), with this being exacerbated in the models of *T ∝ V* and constant volume. This is as a result of the Poisson effect, where an increased demand on conserving the volume when stretched in one direction causes the wall to pull in in the other direction. The relative volume change decreased from constant turgor, to *T ∝ V* and fell to 1 as expected for constant volume (Fig. 5k). This behaviour is not dependent on the cell geometry, which means this type of quantification could give clues as to how plant tissues’ turgor behaves during deformation.

We quantified the area change of the different species during deformation (Fig. 5j) and saw that deformation in any direction of the tissue resulted in an increase in cell area in both Arabidopsis hypocotyls and Maize leaves, which increased significantly (*p <* 0.001 see section 9) as we increased the tissue strain with the response changing with species (*p <* 0.001), however, deformation direction was not significant (*p* = 0.466). Despite variability among samples, the mean response of biological replicates consistently showed a cell area increase. The area increase across Arabidopsis hypocotyls and Maize leaf primordia suggests that stretching induces a common mechanical response in water uptake. This is consistent with the model of constant turgor. On the other hand, *Tradescantia* volume change did not alter and remained on average just above 1, showing its behaviour to be closer to the constant volume formulation. Note that this could also be influenced by the experimental timescale, where the time taken to capture the image could provide time for water uptake.

Comparison with simulations, therefore, suggests the possibility of water movement during deformation and that this behaviour varies between species, thereby altering the tissue response.

### 2.8 Arabidopsis cell division mutant *trm678* has altered tissue properties

We next tested the effect of altered cell division. We compared wild-type Arabidopsis hypocotyls to hypocotyls from *trm678* mutants [Schaefer et al., 2017]. The loss of precision in cell division orientation resulted in the longitudinal walls having a slightly reduced edge extension (Fig. 5c) for smaller tissue deformations but were comparable to the wild type for higher strains. This is explained by the edge deflection, where *trm678* exhibited an increased level of edge deflection (Fig. 5h). To simulate *trm678* we took the average Arabidopsis cell shape and added noise to their junctions (creating five different meshes) (Fig. 5e). These altered outlines when simulated showed agreement with the data, where the deformation of their aligned edges (when deformed in the longitudinal direction) was between Maize and Arabidopsis (Fig. 5f). The model failed to fully capture the edge deflection response (Fig. 5i). This demonstrates that in the longitudinal direction, the altered cell division does little to impact their mechanical response, possibly because, as in *trm678*, the cell arrays are moderately maintained (Fig. 1d). However, examining their response transversely (not possible experimentally)(Fig. 5g), we saw their edge extension was lower than all of the other species, implying this mutant could be more vulnerable to transversal deformations. Consistent with this prediction, we observed that *trm678* hypocotyls markedly differed, with a mean hypocotyl width/diameter of 301.0 ± 31.5 SD compared to the wild type, with a mean width of 202.9*µm* 25.0 SD, which is approximately 48% greater than wild type. Tissue response to deformation can, therefore, be affected by division patterns dependent on the degree and direction of the division plane

## 3 Discussion

### 3.1 Cell topology as a key determinant of tissue mechanics

While cell wall composition is undeniably central for regulating cell wall mechanical properties and those at the tissue level, our findings reveal a significant geometric layer of regulation. By controlling for cell density and wall properties, and integrating 3D mechanical simulations of turgid cells with quantitative live-tissue experiments, we demonstrated that deformability is tunable purely through cell shape and junction arrangement. This implies that cell geometry is critical in defining the mechanical state of a tissue and determines tissue resilience and strain accommodation, and is not just dependent on the cellular density [Malek and Gibson, 2017, 2015] or the plant cell wall properties themselves [Smithers et al., 2024, Cosgrove, 2005, Peaucelle et al., 2012, Hamant and Traas, 2010]. These properties are critical to how plants respond to deformation. Our work builds upon extensive investigations into plant cells as cellular solids [Gibson, 2012]. Our work is particularly relevant during early development, when primary walls lack the reinforcement of secondary cell walls [Cosgrove and Jarvis, 2012]. Our data suggests that geometric tuning allows tissues to achieve mechanical robustness at a low metabolic cost. We also revealed the consequences of different cellular geometries on the distribution of maximal strain. By optimising the redistribution of internal strains, plants can maximise their size or resilience using minimal biomass, providing a clear competitive advantage in environments defined by unpredictable mechanical perturbations, such as wind or gravity [Moulia et al., 2021].

We examined the impact of altering the junction-to-junction distance, internal angle and edge length ratios. We observed that four-way junctions have higher strain upon deformation, which could potentially explain why, when creating air spaces, certain plants (for example, rice and maize roots in transverse section) [Sinnott and Bloch, 1941] divide to form four-way junctions between distinct layers of root cortical cells. Consistent with previous observations [Erguvan et al., 2025], three-way junctions displayed a mechanical strain hotspot. This would suggest that four-way junction avoidance is not an attempt to minimise strain hotspots.

We identify three-way junctions as a fundamental mechanical element that functions as a hinge, allowing for junction angle uniformity changes that efficiently redistribute strain during deformation. In contrast, the relative stiffness of four-way junction setups restricts this mode of deformation, suggesting that the observed biological avoidance of four-way junctions [Gascon et al., 2025, Wang et al., 2024b] may be an evolutionary strategy to optimise mechanical flexibility.

The flexibility of the three-way junction is further enhanced by arranging the walls to create perfect 120*^◦^* junctions. Creating hexagonal three-way junction arrays may be a natural phenomenon where junction angles commonly evolve to be 120*^◦^* [Wang et al., 2024b, Lloyd, 1991, Korn, 1980, Bonfanti et al., 2023]. Junction uniformity is thought to be the product of the three walls’ tension equalising, as in bubble foams [Wang et al., 2024b, Lloyd, 1991, Flanders et al., 1990].However, the selective stiffening of new cell walls in some species can cause junctions to converge to 120*^◦^* more quickly during growth, accelerateing the formation of a more flexible structure [Bonfanti et al., 2023]. The placement of the new division wall is also crucially important as demonstrated by the analysis of the *trm678* mutant. While the cells of the leaves of Maize versus *Tradescantia* had quite different responses to deformation when division plane orientation was slightly perturbed in the *trm678* mutant the balance between junction angle change and cell wall strain was altered by a similarly small amount, suggesting fine-tuning of the properties is possible.

Further work is needed to fully understand the mechanism by which different cell shapes are created. The possible advantage of three-way junctions, combined with evidence of four-way junction avoidance Gascon et al. [2025], Wang et al. [2024b], Martinez et al. [2018], suggests that this now needs to be an extra consideration of the extent to which a four-way junction is avoided, as close three-way junctions (within 10%) behave similarly to four-way junctions. Avoidance that we see in actual plants by a margin of around 10% of the edge length Gascon et al. [2025], Sinnott and Bloch [1941], thus providing a quantitative parameter for four-way junction avoidance in future implementations of division rules [Serra and Robinson, 2020, Besson and Dumais, 2011, Bouchez et al., 2024, Müller, 2012, Goldy et al., 2026].

Cells with longer edges experience greater deflection for the same junction change, implying that rectangular setups could be advantageous over square ones for the same surface area in reducing wall strain. Additionally, our results have revealed a possible conflict between larger cells with longer edges having reduced strain upon deformation and the impact of turgor stress [Sapala et al., 2018] and epidermal tension [Silveira et al., 2025]. To combat this conflict, plants may use lobed pavement cells which minimise turgor stress [Sapala et al., 2018] and toughen the plant [Bidhendi et al., 2023], demonstrating that there are more shapes to explore than the simple ones appearing in this article.

### 3.2 Internal layers and strain hotpots

Previous literature surrounding this topic has largely focused on epidermal or single-cell-layer deformations. The role of internal layers in morphogenesis is increasingly being recognised [Silveira et al., 2025, Mosca et al., 2017, Yates et al., 2026]. Our results demonstrate that the impact of different cell geometries is altered in a 3D tissue context compared to a monolayer. The cortex cells also exhibited different strain behaviour upon deformation (i.e., a greater increase in strain), further underscoring the need to consider the inner layers. Strain hotspots, inefficient dissipation and redirection of tissue stresses induced by turgor and external load, require plants to invest in thicker cell walls, resulting in higher resource allocation costs [Bidhendi and Geitmann, 2016, Jędrzejuk and Kuźma, 2025]. Patterns of mechanical stress within these cellular networks feed back onto growth and development, influencing microtubule alignment, cellulose deposition, and division orientation [Hamant et al., 2008, Robinson and Kuhlemeier, 2018] thus, poor management of tissue stresses could affect development. Understanding the distribution of stresses upon deformation is important as we work to understand how plants respond to mechanical perturbations.

### 3.3 Water movement

Our results show that the way in which water is modelled can affect the strain on the walls, especially whether orthogonal walls are relatively compressed, as well as the overall material properties. In a previous study when comparing constant volume and constant turgor, only minor changes in the response to compression were observed [Weber et al., 2015]. Additionally, allowing a drop in turgor pressure [Diels et al., 2019], assuming a permeable membrane [Forterre and Jensen, 2022], also made little difference to the data fit. However, these compression experiments were completed under a short timescale and in a single cell. It has been shown that having higher or lower turgor can significantly change the behaviour of the cells. At high turgor pressure, taut cell walls deform immediately through stretching, whereas in low-turgor walls initially deform by bending before matching the same behaviour of high turgor. [Gibson, 2012]. By comparing model predictions with experimental data our work shows that water movement during deformation might vary between species and organs. There is emerging work on measuring water conductivity and behaviour in the tissue context of *Marchantia* [Laplaud et al., 2024], where other models have examined water movement in 2D on the growth timescale [Cheddadi et al., 2019, Long et al., 2020, Oliveri et al., 2026]. Further study is needed to better understand water movement during deformation.

### 3.4 Next steps in mechanical modelling

While our model is able to capture much of the behaviour we see in the experimental data, future work can enhance mechanical fidelity by transitioning to 3D solid models (removing the thin-film assumption of the cell walls) [Malek and Gibson, 2017, Shafayet Zamil et al., 2017, Gascon et al., 2025, Majda et al., 2017, Lee et al., 2025], enabling direct access to compressive regimes, shear stresses, and through-thickness stress gradients [Malek and Gibson, 2015, 2017]. Alternatively, shell formulations could be extended to incorporate rotational forces at junctions, capturing more detailed junctional bending and twisting behaviour [Jensen and Revell, 2023]. To balance computational efficiency with structural detail, future studies could adopt a multiscale approach—leveraging fine-grained solid models at the cellular scale to parameterise coarse-grained tissue-scale simulations [Saikia et al., 2020, Shafayet Zamil et al., 2017, Ghysels et al., 2009, Boudaoud et al., 2023]. Incorporating structural anisotropy to account for directional cellulose alignment and accurate models of water movement will further refine stress-strain predictions [Lee et al., 2025, Chen et al., 2025].

## 4 Conclusion

Mechanics is not the only consideration constraining plant cell shape, size and arrangement. There are accommodations to maximise photosynthesis [Egesa et al., 2024], organ function [Kierzkowski and Routier-Kierzkowska, 2019], cellular connectivity [Carter et al., 2017]. or to produce specialised cells [Mathur, 2004, Whitney et al., 2009, Woolfenden et al., 2018, Bergmann and Sack, 2007, Liu et al., 2021, van Spoordonk et al., 2023, Meyer et al., 2017]. All of which will constrain cell geometry and ultimately impact the mechanical properties of the plant.

These findings extend beyond plant biology, offering insights for tissue engineering and materials science. The ability to tune the mechanical properties of a tissue or a material by adjusting its vertex connectivity or cellular anisotropy independent of the material’s intrinsic composition provides a framework for designing tunable biomaterials. Just as plants utilise geometric cellular configurations like hexagonal arrays to create soft and flexible structures, synthetic porous scaffolds or bone implants could be optimised by incorporating similar junctional hinge mechanisms to improve strain distribution and prevent fatigue failure [Marin, 2023, Pais et al., 2023, Wegst et al., 2015]

In summary, this work provides a quantitative link between tissue architecture and mechanical resilience. We have shown that cell division patterns—and the resulting junctional topology—are not merely passive consequences of growth, but are active tuning mechanisms. By avoiding energetically costly four-way junctions and adopting efficient packing geometries, plants achieve a dynamic range of mechanical performance, allowing them to adapt to diverse developmental and environmental constraints. Although deformation mechanics intuitively link to organ expansion, fully deciphering how cellular geometry and altered strain patterns govern this growth requires additional computational modelling and experimental validation.

## 5 Acknowledgments

The authors would like to thank Dora Cano Ramirez for gifting us a cutting of the *Tradescantia* used in the experiments. We thank Magalie Uyttewaal for the kind gift of the *trm678* X pUBQ10::29-1tdTomato seeds. We would also like to say a big thank you to the Cambridge University Herbarium for helping us identify and advise on the imaging, and for granting access to the samples shown in Fig. 6, especially Anne Dubearnes, Amber Horning, and Juliet Anderson, in no particular order. Thanks also go to Chiara Perico and the Langdale lab, who provided excellent advice on caring for *Zea mays* and dissecting them. We would also like to thank the rest of the Robinson group and the whole of the Whitewood and Schiessl teams for the great feedback during lab meetings. We are grateful to the professional services at SLCU for all their help, particularly Horticulture and Microscopy. We thank Lauren Wier for plant care.

## 6 Author contributions

S.R. supervised the project. S.R and L.S Project conception and preliminary data. E.S wrote the original draft with editing by S.R and M.E. Feedback was provided by all authors. E.S built the simulations tool and ran the simulations. M.E designed and performed all the experiments and segmented the images. E.S completed the analysis of these images with neighbour extraction code provided by E.L. M.L wrote the updated code for the ACME experiments.

## 9 Methods

Here we detail the interdisciplinary approach and methods in this paper. The computational methods are detailed in section 9.1 and the experimental methods in section 9.2.

### 9.1 Computational methods

#### 9.1.1 Shape fitting with set perimeter and area

Here, we set up the equations used in Fig. 2 to understand the optimum shape to fill a tissue regarding stress induced from turgor pressure. This work assumes that each cell uses a defined amount of resources for its parameter (anticlinal wall) and surface area (periclinal wall), where we are not considering any boundaries of the tissue domain (as taking a sample from an infinite domain). Starting with hexagons, let *l_H_* be the length of the hexagon’s sides, *P_H_* their individual perimeter, *S_H_* the surface area, then *TA_H_* the total area (including both the bottom and top of the cells), and *TP_H_* the total perimeter. The perimeter is,

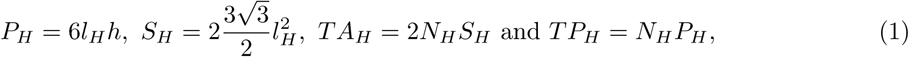

where *N_H_* is the total hexagon cell number we need to find and *h* the height of the walls. Likewise, for squares with side length *l_S_*, *P_S_* the individual perimeter, and *S_S_* the surface area, the total area *TA_S_*, and perimeter *TP_S_* is,

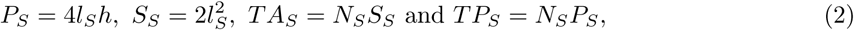

where *N_S_* is the square cell number. Finally, for the rectangles which have one side of length *l_R_* and the other with length *αl_R_*, where *α >* 1 is the aspect ratio, then their individual perimeter *P_R_*, surface area *S_R_*, and total area *TA_R_*, perimeter *TP_R_* are,

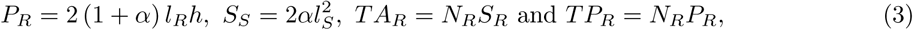

where *N_S_* is the rectangle cell number. We fix the total surface area and perimeter, and rearrange the equations for the cell numbers to get,

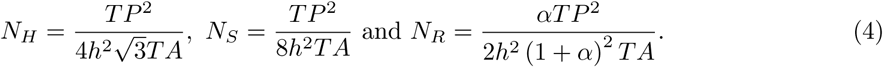

From this, we can rearrange the surface area equations to get the cell lengths,

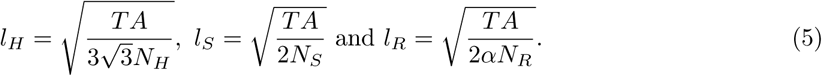

With these lengths, we now find the radius of the maximum circle, *r_H_*, *r_S_* and *r_R_* that fit in a hexagon, square and rectangle, respectively, which are,

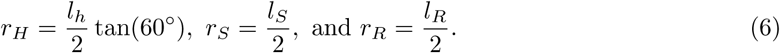

#### 9.1.2 Finite element methods

To determine the impact of geometry on the mechanical properties of plant tissues, triangular meshes with a Delaunay-based unstructured mesh generation algorithm from Darren Engwirda [Engwirda, 2014] were generated of different cellular solids of idealised plant tissues in 3D. All meshes were created so that the triangle edges on the cell boundaries were approximately 0.5*µm* apart. The meshes were built using custom MATLAB code (set up to facilitate easy use) that can create a range of different 3D setups with varying numbers of layers and hypocotyl-like structures (available at https://gitlab.developers.cam.ac.uk/slcu/teamsr/meshing_code.

For the monolayer structures, we create square staggered (three-way junctions) with a bigger variant (15*µm* in edge length instead of 10*µm*) and non-staggered (four-way junctions), rectangular staggered (three-way junctions) and non-staggered (four-way junctions) with rotated variants, and hexagon 3D meshes. Meshes of hexagons with varying internal angles, squares of varying staggering and the average cell shapes for each species were also made. All these meshes have the same top surface area.

For multilayer tissues, we began with three layers of square and rectangular cells, with a cortex sandwiched between two epidermal layers. In these meshes three-way and four-way junction variants were created with smaller epidermal cells (epidermal cells and cortex cells being 10*µm* and 15*µm* in height and width respectively) where we then created additional 3 way junction meshes with varied cell size in the layers, specifically, one with bigger epidermal cells (epidermal cells and cortex cells being 15*µm* and 10*µm* in height and width respectively) and one with even layers (all layers 10*µm* in height and width).

Lastly, a hypocotyl cylindrical structure was built with 4 central cells and 20 epidermal cells, with non-staggard, staggard, and hexagonal variants to create a realistic plant structure, all with the same height and radius.

All these meshes was then converted into a compatible form to inflate in the Tissue software originally from the Jonsson group [Hamant et al., 2008, Bhatia et al., 2016, Bozorg et al., 2014, Bonfanti et al., 2023], which has now been further developed and optimised (available at https://gitlab.com/slcu/teamHJ/tissue/-/tree/petsc?ref_type=heads in the PETSC branch). Tissue uses a finite element method approach using the discretised mesh surface to approximate solutions to the hyperelastic continuous mechanical equations, which are being stretched via an applied pressure force in each mesh triangle. It uses the ‘Triangular Bi-Quadratic Springs’ (TRSB) method, which uses a St Venant Kirchhoff formulation and approximates it using biquadratic springs, which resist triangle edge and inner angle deformation [Delingette, 2008] and allows us to simulate cell walls efficiently. For further details see section 10.1. Mesh convergence studies for our solver and cellular meshes were already completed for previous publications [Bonfanti et al., 2023]

Using the TRSB method, a stress tensor is calculated for each triangle, where the maximum stress and strain magnitude and direction are found from these tensors from the maximum corresponding eigenvector and eigenvalue. The average strain per simulation *s̄* is calculated using the weighted average of the maximum strain magnitude on each triangle in the centre cells weighted by the triangle area. For the anisotropy implementation used in Fig. 7f, see [Bozorg et al., 2014] for details, where we have no stress feedback and instead set the anisotropy direction in the periclinal to be against the cell file, with no anisotropy contribution in the anticlinal wall.

The pressure force is applied to all the triangles proportional to their area in the outward normal direction. A solution to the simulation is found when the system has reached mechanical equilibrium, i.e., when the pressure forces acting on each node from its surrounding triangles are balanced by the strained triangles surrounding it, so that the total force is 0. We find this mechanical equilibrium using the Newton-Raphson method [Press et al., 2007, Amrein and Hilber, 2020, Amrein and Wihler, 2014] with adaptive time stepping and a tolerance of 10*^−^*^5^ (and 10*^−^*^3^ for the hypocotyl meshes).

All simulations were inflated with a turgor pressure of 1MPa, Young’s modulus of 100MPa and Poisson ratio of 0.4 in line with values set in the literature [Bonfanti et al., 2023, Majda et al., 2022]. We have implemented three different methods of handling the turgor pressure. For constant turgor, the same turgor value is used throughout the simulation. For *T α V*, we assume water behaves like a gas, which assumes that the deformation does not increase temperature (dispelled quickly) [Vesenjak et al., 2007]. Thus, for a cell *i* the turgor pressure is,

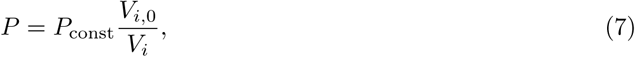

where *P*_const_ is the set constant turgor pressure *V_i_*, the current volume of the cell and *V_i,_*_0_, the initial volume of the cell. For constant volume we iterate the turgor pressure value by making small decreasing/increasing steps to it value proportional to 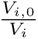 until we reach a level of tolerance (an average of less 5*µm*^3^ difference between *V_i,_*_0_ and *V_i_*)

For the set deformation of the meshes, we first inflated them and then applied a linear transform to the coordinates (multiplying the coordinates by 1.2 in the direction of deformation). All coordinates on the stretched direction boundary were then fixed in the direction of the stretch with Dirichlet boundary conditions [Granzow, 2023]. To deform the meshes with a set force, we applied forces to each boundary node uniformly in the deformation direction, with the total amount being the same for each simulation set (total force per side for the hypocolty simulations being 24720 and for the square and rectangle monolayer 10880).

We therefore control for mesh density, stiffness, and turgor pressure. The only variable is then the shape of the cells. All analysis and data are performed on the central cells to exclude any boundary effects.

### 9.2 Experimental methods

#### 9.2.1 Plant growth conditions

*Zea mays* plants were grown in 5cm square pots for no longer than 2 weeks. Growth conditions were set as 16 hour/8 hour day/night cycle, temperature 28*^◦^C/*20*^◦^C*, and light intensity of 250*µ*mol m*^−^*^2^s*^−^*^1^ . They were planted in a *Zea mays* mix of John Innes No.2: Levington’s M3: Vermiculite in a 2 : 1 : 1 ratio. Seeds were sourced from the Rasmussen lab [Martinez et al., 2017].

*Tradescantia zebrina* were potted in 9*cm* square pots in John Innes Number 2. They were grown in a greenhouse at the Sainsbury Laboratory, University of Cambridge, United Kingdom. Details of these conditions are 16-hour days with supplementary light (min radiance 88*wm^−^*^2^), day/night temperature approximately 25*^◦^C/*15*^◦^C*, with cooling occurring on 30*^◦^C* and shading closed at 500*wm^−^*^2^. *Tradescantia zebrina* was sourced from a garden centre.

Arabidopsis plasma membrane marker line pPDF1::mCitrine-1xKa1 (PM-YFP) was used as previously described [Simon et al., 2016] and *trm678* × pUBQ10::29-1tdTomato [Melogno et al., 2024]. Arabidopsis seedlings were surface sterilised with 5% bleach and were grown on vertically placed 90*mm* Petri dishes in 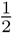 MS media for 7 − 10 days. Plants were grown in long day conditions (LD) (16 h light/8 h dark) with a photosynthetic photon flux density (PPFD) of 170 *µmol/m*^2^*s*.

#### 9.2.2 Image acquisition for cell geometry quantification

The live 3D images of Arabidopsis, Maize and *Tradescantia*in Fig. 1 were acquired using Leica Stellaris 5 confocal microscope using a multi immersion objective (×25, 0.95) in water immersion mode. For *Tradescantia* imaging, the youngest half opened leaf was taken to peel epidermis from the middle part. For quantification in Fig. 1, live 3D stacks of Arabidopsis, Maize, and *Tradescantia* were acquired for downstream analysis. Arabidopsis and Maize were imaged using a Leica SP8 confocal microscope (Leica Microsystems) with a PMT detector and a HCX APO L 20 × */*0.5 W objective. *Tradescantia* was imaged using a Leica Stellaris confocal microscope (Leica Microsystems) equipped with a HyS detector and a multi-immersion HCPL-APO CS2 20 × */*0.75 objective in water mode. All acquisitions used LAS X software. For Arabidopsis, plasma membrane (PM-YFP) was excited with an OPSL 514*nm* laser (3%) and detected between 519 − 557*nm*, with PMT gain of 32V. Maize cell walls were stained with Calcofluor White (Fluorescent Brightener 28 disodium salt), excited with a diode 405*nm* laser (laser power: 20%) and detected between 439 − 529*nm*, with PMT gain of 496V. The youngest half-opened leaf of *Tradescantia* was selected to peel from the middle part of the leaf for imaging. Cell wall autofluorescence of *Tradescantia* was excited with a diode 405*nm* laser (laser power: 20%) and detected between 426 − 517*nm*, with HyS detector gain of 6.8%. All images were acquired using bidirectional scanning at 400Hz with no frame or line averaging and z thickness of 0.5*µm*.

The moss samples in Fig. 6 were selected from microscope slides held in the Cambridge University Herbarium. Images of these specimens were acquired using a VHX-7000 digital microscope (Keyence, Osaka Japan).

#### 9.2.3 Extensometer experiments

A modified version of the ACME was used to apply uniaxial deformation to the samples. The hardware components have been updated to match current availability and to take advantage of the advances in the hardware. Load cell FSH03868 - LSB200-20g, Jr S-Beam Load Cell as used previously, with FSH04720 – USB225, High Resolution USB Output Kit and SLB00006 – Integration, Configuration & Calibration Load Cell with USB kit NIST Traceable Certificate (Futek) enables the load cells to be plugged directly to a computer without the need for amplifiers or additional calibration steps. The new setup is therefore easier to assemble and more portable. The positioners are as used previously SLC-1730-D-S Positioner with 21 mm travel with an updated controller MCS2-S-0001 MCS2 Sensor Module and MCS2-C-0001 MCS2 control system (Smaract). A new version of the software was also developed to match the new components https://doi.org/10.5281/zenodo.21372723.

Samples were prepared and attached to ACME plates as follows. The *Zea mays* plants were then dissected when they were at least a 2 week old to extract a young leaf near the primordia that was approximately 2cm long. Samples were stained with a 2.5% solution of Fluorescent Brightener 28 disodium salt (Sigma-Aldrich, 4193 55 9 50 mL) for 20 minutes and subsequently washed with ddH2O three times. Leaf primordia were cut into 0.5 − 1*mm* thin sections (measured from the image) longitudinally for axial stretch and transversely for transverse stretch and uncurled using a needle (AGANI*^T M^* 0.3 13*mm*) at the time of mounting on tough tags as in Robinson et al. [2017] and glued using superglue (Loctite Precision Super Glue). The top half-opened leaf was cut from the tip of a branch. The abaxial epidermis was peeled from the upper part of the leaves (as in supplementary) using a needle, cut into 0.5 − 1*mm* in both longitudinal and transverse directions. Before stretching, images were acquired using a Leica SP8 with a HCX APO L 20 × */*0.5-W objective and a Leica PMT detector. For Maize imaging, leaves stained for 30min with 2.5% working solution of Fluorescent Brightener 28 disodium salt solution (Sigma-Aldrich 4193 55 9) were excited using a diode 405*nm* laser and detected using a PMT detector at 439 − 529*nm* with laser power set to 20% with gain set to 496 V. *Tradescantia* epidermal cells were imaged using a diode 405 nm laser at 20% and autofluorescence was captured using PMT detector at 421 − 611*nm* with gain set to 842 V using bidirectional scanning at 400 Hz. Arabidopsis control plasma membrane marker (PM-YFP) was excited with an OPSL 514nm laser (3%) and detected between 519 − 557*nm*, with PMT gain of 32V. The *trm678* × pUBQ10::29-1tdTomato line was imaged using OPSL552 laser (2%) with HyD2 detector at 557 − 605*nm* at gain 19V. To avoid tissue slippage, force was applied in increments of 0.5, 1, and 2 g for Maize, and up to 8 g for Tradescantia until roughly 20% deformation was achieved. After stretching, images were collected as soon as possible to keep stress relaxation to a minimum by keeping the acquisition time under 5 minutes, depending on sample z-thickness. All images were acquired using a z thickness of 0.5*µ*m, with a scan speed of 400 Hz and without line or frame averaging.

#### 9.2.4 Cell segmentation

Maize confocal image stacks were processed using MophographX [Strauss et al., 2022]. Briefly, cell outlines were extracted from the plasma membrane signal using a 2.5D segmentation pipeline. A curved mesh representing the epidermal surface was generated by applying Gaussian blur of 0.3 − 0.7 and edge detect between 2000 − 5000, marching cube of size 5 after which the intensity of the signal was projected onto the mesh. Mesh was divided and smoothed 2 − 3 times and individual cells were segmented using watershed segmentation. Segmentation quality was checked visually and corrected manually where needed to ensure accurate cell boundaries.

*Tradescantia* epidermal cell boundaries were segmented from 2D maximum intensity projections of confocal images using Cellpose (V3.x) [Stringer et al., 2021], using the pretrained Cyto2 model. Automated segmentation masks were generated using default model parameters and were visually inspected for accuracy. Where necessary, segmentation errors were corrected manually. The resulting segmentation masks were exported for downstream analysis.

The contours are extracted from these masks. These contours contained segmentation errors that needed to be corrected. For this, we used custom code that allowed us to manually move, delete nodes and create new edges (available on GitHub https://gitlab.developers.cam.ac.uk/slcu/teamsr/edit-2d-segmentation). With the corrected points and edge information, we then identified each cell (finding the sets of edge indices that lie in a closed loop by looping through edges and going clockwise at each junction). To pair up the cells before and after we took advantage of the unchanged cell patterning during the tissue stretching, we used the cell agency graph to match the segmentation before stretching and after stretching. From one couple of cell labels before and after the stretch, we iteratively spread the mapping along the graph by optimizing the local match of topological cell features (percentage of contact, relative orientation etc). The optimisation of the local graph matching is achieved using the pygmtool python package [Wang et al., 2024a]. All this processing code was written in Python and can be found on GitHub (https://github.com/L-EL/graphMatching).

#### 9.2.5 Cell shape and edge analysis

To examine edge strain and enable comparison with simulations, we characterise each edge into four categories using a set of criteria. We first label each edge aligned and non-aligned if its direction is within 45*^◦^* of the stretch direction. We then label edges Longitudinally stretched, aligned walls (if they are aligned and connected to one non aligned wall and one aligned wall at each end), Longitudinally stretched, non-aligned walls (if they are non aligned and connected to two aligned walls at each end), transversely stretched, aligned walls (if they are aligned and connected to two non aligned at each end), transversely stretched, non aligned walls (if they are non aligned and connected to one non aligned wall and one aligned wall at each end).

We calculated the junction-to-junction distance % for each junction, by identifying the longitudinal edges (see section 9 for criteria), and using these two edge lengths (*l*_1_ and *l*_2_ Fig. 1i) to get the percentage (min(*l*_1_, *l*_2_)/*l*_1_ + *l*_2_).

The edge deflection is calculated by taking each edge vector’s component (normalised) in the stretch direction (the dot product) after deformation and finding the ratio of the projected distances from the edge length before deformation divided by the after. This measures how much deformation in the stretch direction you get from just the edge reorienting, so the closer you are to 1, the less the edge has deformed.

To extract the internal angles, we first identified each cell that has 6 sides. Each cell was then identified as being in the cell file direction or against it. For the *Tradescantia*, this was found by calculating the vectors from the centre point of the current cell to all its neighbours, and the midpoint vectors, which were the average direction between two neighbours next to each other. We then examined the difference in angle between the stretch direction of the tissue and pairs of neighbour vectors and pairs of midpoint vectors, where each pair consists of vectors on opposite sides of the cell. The minimum of all pairs’ total angle differences with the strain direction for the neighbourhood and the midpoint direction was found. If the minimum angle difference from the neighbour vectors was less than the midpoint difference, then the cell was considered to be in the cell file direction and vice versa. As *Zea mays* and Arabidopsis Wild type and *trm678* were in strict columns, each cell was labelled to be in the cell file or against it if the specimen was deformed with or against the cell file. Internal angles were calculated from the edge vectors (junction-to-junction) in each hexagon. For cells in the cell file direction, we selected the four edges with the smallest angles relative to the stretch direction, and for cells against the cell file, we selected the four largest angles. We then use the average angle these four angles make with one another when they are connected, calling it *ϕ* (i.e., there will be two pairs and two angles). The internal angle depicted in Fig. 1 is then 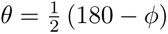.

We calculated the tissue strain from the images using the junctions as nodes labelled **X** and **x** for the before and after deformation positions in a finite element mesh. These nodes were then triangulated using MATLAB’s Delaunay triangulation algorithm. We then found the deformation gradient 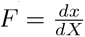, the right stretch tensor *U*, (where *F* = *RU*) and then linear the Biot strain tensor *E* = (*U* - *I*) where *I* is the identity matrix. The strain was then extracted in the stretch direction, and the tissue deformation was set as the weighted-average (triangle-area) strain.

For all analyses of the experimental stretching data, we conducted ANOVA to assess significant relationships between the variables and edge strain. Specifically, Fig. 5c-d revealed that all edge Direction (*p <* 0.001), species (*p <* 0.001) and stretch direction (*p <* 0.001) significantly determined the variance in edge strain response. Crucially, a significant Species × edge direction interaction ( *p <* 0.001) demonstrates that the directional mechanical response of cell edges varies depending on the species. A linear model incorporating just Maize and Tradescantia (allowing us to examine stretch direction in detail) and performing an ANOVA showed the same significant relationships; furthermore, a significant stretch direction × edge direction interaction (*p* = 0.039) indicates that the deformation direction changes edge-level strain. For Fig. 5h, an ANOVA revealed that stretch direction (*p <* 0.001), tissue Strain (*p <* 0.001) and species (*p <* 0.001) all significantly drive the response. Crucially, two-way interaction terms confirmed that the rate edge deflection across tissue strain varies depending on the stretch direction (tissue Strain × stretch Direction: *p <* 0.001) and species (tissue Strain × species: *p <* 0.001). A significant three-way interaction (tissue Strain species stretch Direction *p* = 0.010) further indicates that species-specific differences in shape change sensitivity are dependent on whether loading is applied transversely or longitudinally. For Fig. 5j, an ANOVA indicated that cell strain was significantly driven by tissue strain (*p <* 0.001) and species (*p <* 0.001), with a significant species-by-tissue-strain interaction (*p <* 0.001). Measurement direction was not significant (*p* = 0.466) and showed no interaction effects (*p >* 0.428 across all terms).

## 10 Supplementary

### 10.1 Finite element mechanical model details

In more detail, the St Venant Kirchhoff formulation stress-strain relation for an isotropic material,

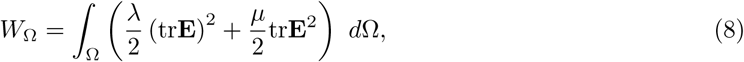

where *W* is total strain energy in a domain Ω, *λ* and *µ* the Lame coefficients and defined in plane elasticity as 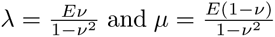 with *E* being the Young’s modulus and *ν* the Poisson coefficient, is approximated over a triangle, *T*, as

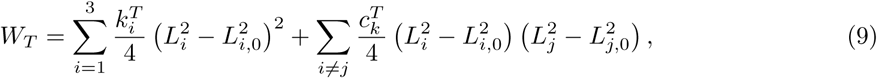

where 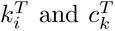 are the tensile and angular stiffness of the biquadratic springs and *L_i_* and *L_i,_*_0_ the current and resting lengths of edge *i* in the triangle. These stiffnesses are defined as

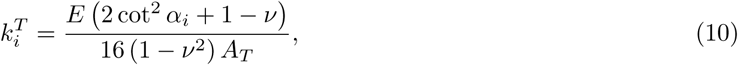

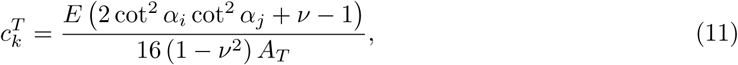

where *α_i_* is the angle opposite edge *i* in the resting triangle configuration and *A_T_* the resting triangle area. A force on each node can then be calculated from this strain energy as a result of the stretching triangles.

### 10.2 Supplementary figures

**Figure 6:**
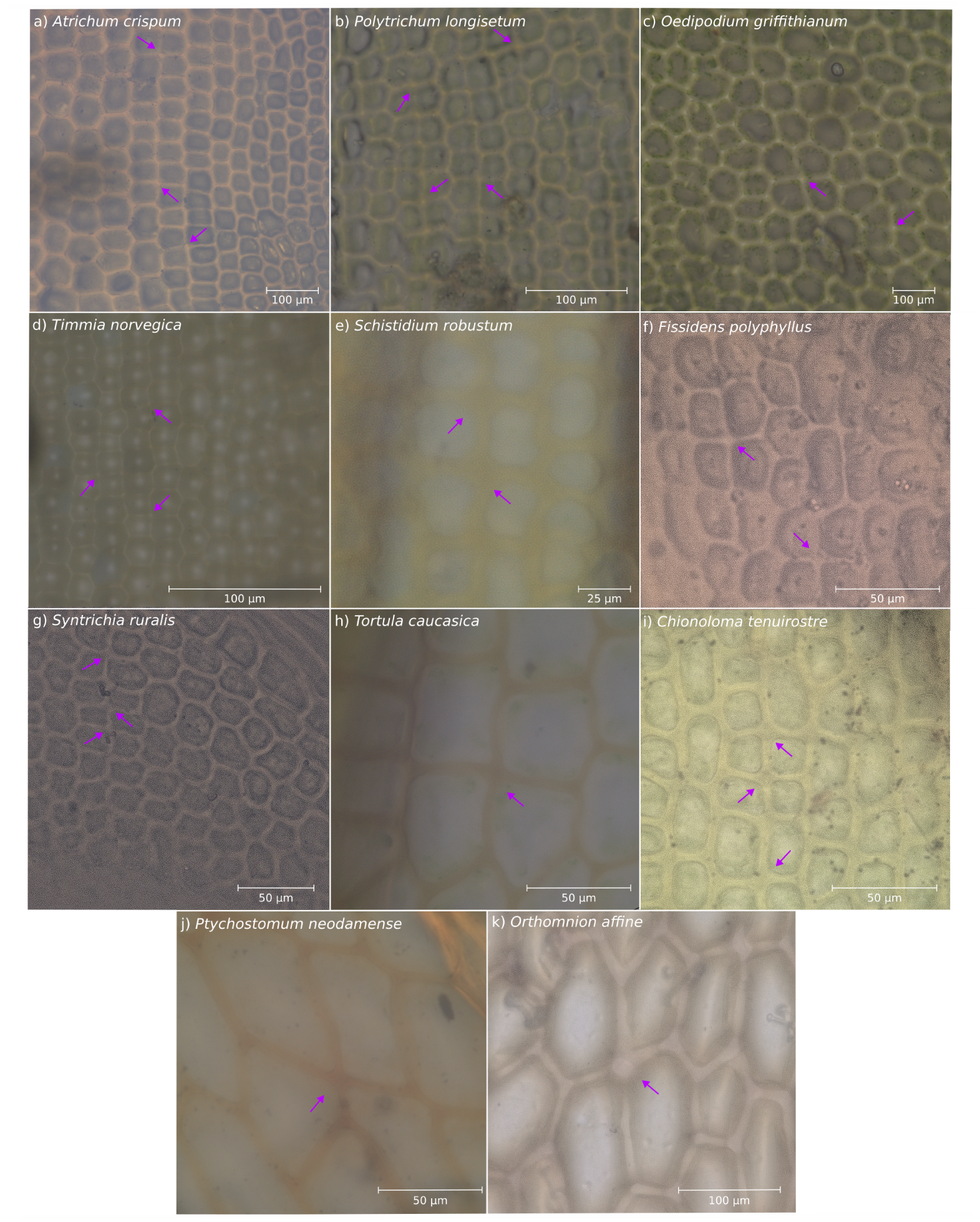
Evidence of four-way junctions (indicated by arrows) across Bryophyte species. Images were taken from slides held at the Herbarium collection ‘Cambridge University Herbarium (CGE)’. a) *Atrichum crispum* b) *Polytrichum longisetum* c) *Oedipodium griffithianum* d) *Timmia norvegica* e) *Schistidium robustum* f) *Fissidens polyphyllus*, g) *Syntrichia ruralis*, h) *Tortula caucasica*, i) *Chionoloma tenuirostre*, j) *Ptychostomum neodamense*, k) *Orthomnion affine*.

**Figure 7:**
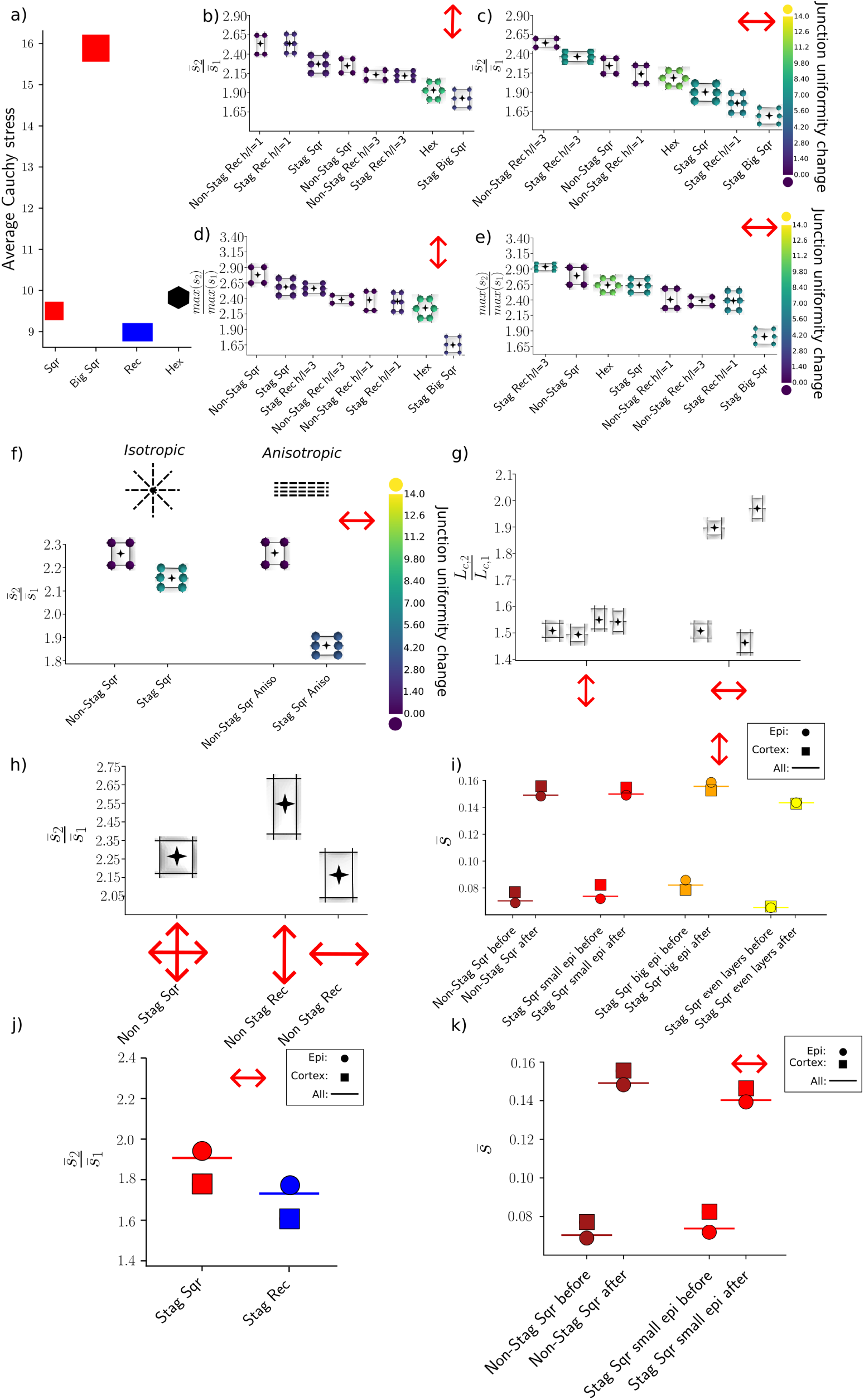
Simulations and quantification of monolayer and multilayer geometries under different conditions provide further evidence that three-way junctions reduce strain upon deformation. a) The average principal Cauchy stress of each mesh triangle on the top surface of a single inflated cell with shapes of a square (of size 10*µm*), a bigger square (15*µm*), a rectangle and a hexagon, all with the same top surface area (apart from the bigger square variant). b-c) The average strain ratio of all the different cell geometries repeated for non-constant turgor when deformed in the b) cell file direction and c) against the cell file with the star indicating the value on the scale. Overlaid are the cellular outlines post-deformation, with dots indicating the junction uniformity change. d-e) The ratio of the maximum strain on the top surface wall before and after deformation of all the different cell geometries when deformed in the d) cell file direction and e) against the cell file, with the star indicating their value on the scale. Overlaid are the outlines post-deformation with dots showing the junction uniformity change. f) The average strain ratio of staggered and non-staggered squares when given isotropic properties (on the left) or anisotropic properties perpendicular to the cell file on their top surface wall (on the right) when deformed perpendicular to the cell file. The star indicates the value on the scale. Overlaid are the outlines post-deformation with dots showing the junction uniformity change. g) The ratio of the average cell length before and after deformation in the deformation direction of staggered and non-staggered squares and rectangles, when stretched with and perpendicular to the cell file with a set force. The star indicates the value on the scale. Overlaid are the outlines. h) The average strain ratio on the monolayer simulations for non-staggered squares and rectangles deformed with and perpendicular to the cell file. The star indicates the value on the scale. Overlaid are the outlines post-deformation. i-k) The average strain/strain ratio in the multilayered simulations with the circles representing the epidermis values, the square the cortex values and the straight line the average of all layers before and after deformation in the i) cell file direction and j-k) perpendicular to the cell for i) non-staggered square, staggered squares with a smaller epidermis, staggered squares with a big epidermis and even layers, j) staggered squares and rectangles and k) non staggered squares and staggered squares.

**Figure 8:**
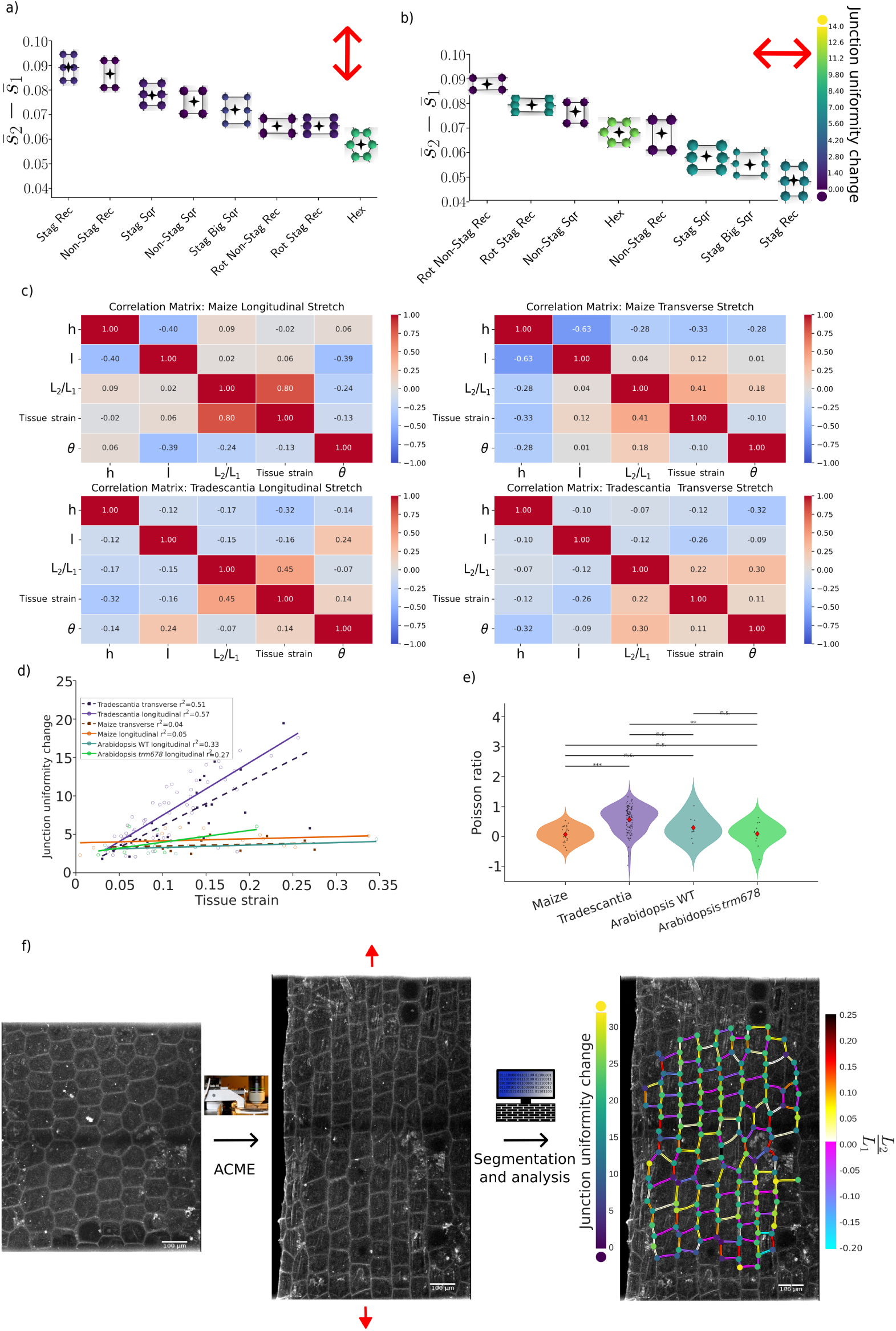
Further quantification of the simulations demonstrate three-way junctions to reduce strain and that live tissue specimen’s deformation response is dependent on several geometric factors. a-b) The average strain difference in the monolayer simulations of all the different cell geometries when deformed in a) cell file direction and b) against the cell file, with the star indicating the value on the scale. Overlaid are the outlines post-deformation with dots showing the junction uniformity change. c) The Spearman correlation value of each cell’s average edge length ratio with the tissue strain of the sample and each cell’s internal angle, deformed with or against the cell file for *Tradescantia zebrina* and *Zea mays*. In total, there are 1924 cells from 100 independent samples of *Tradescantia zebrina* and 928 cells from 27 independent samples of *Zea mays*. d) The junction uniformity change against tissue strain from the experimental data. e) The Poisson ratio from the experimental data using Welch’s ANOVA (*p <* 0.0001) and a post hoc Games-Howell test. f) Diagram showing stretching procedure and analysis. On the left, a confocal image of *Tradescantia* specimen before stretching in its relaxed but turgid state. The specimen is then deformed (red arrows) using ACME (automated confocal micro-extensometer), as shown in the middle. Both images are segmented and mapped onto each other, allowing calculation (image on the right) of the junction uniformity change (dots at the junctions) and the edge length ratio 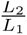 (coloured edges).

